# On the causes of correlated genomic ancestry across contrasting hybridization histories in a monkeyflower species pair

**DOI:** 10.64898/2026.04.02.716215

**Authors:** Matthew C. Farnitano, V. Alex Sotola, Andrea L. Sweigart

## Abstract

Hybridization is a powerful force shaping the evolutionary trajectories of species, yet its outcomes are highly variable both across taxa and within a pair of species. In this study, we examine the processes shaping variation in the extent of hybrid ancestry, both among populations and across the genome. We use low-coverage sequencing data to infer local ancestry across the genome for 782 individuals from multiple populations across two geographic regions within the broadly overlapping range of *Mimulus guttatus* and *Mimulus nasutus*. We find that the extent of hybrid ancestry is variable across populations, supporting disparate historical patterns of hybridization. However, genomic patterns of hybrid ancestry are correlated across groups, indicating they are shaped by parallel processes. Correlations are highest in geographically proximal populations, including between sympatric and allopatric locations, providing evidence that introgression is not locally constrained but spreads via migration across the landscape. We find that features of the genome are predictive of hybrid ancestry and its correlations among populations. However, contrary to findings in some other species, these patterns are likely not driven by simple linked selection against hybrid ancestry. Genomic outliers for high hybrid ancestry are often shared among populations, suggesting a role for parallel positive selection on ancestry. However, known loci associated with reproductive isolation are poor predictors of ancestry variation across populations, indicating that selection acting in natural hybrid populations is highly polygenic and that the underlying genetic architecture varies across space. Overall, this study demonstrates how ecological, demographic, and genomic features all interact to shape the outcomes of hybridization.

## INTRODUCTION

Hybridization between related lineages is a widespread phenomenon across the tree of life (Abbott et al., 2013; Lotsy, 1931; Mallet, 2005; Stebbins, 1959), and may be increasing as a result of anthropogenic environmental change (Chunco, 2014; Gilman & Behm, 2011; Moran et al., 2025). Patterns of hybrid ancestry (introgression) across the genome can reveal the historical and selective processes shaping hybridization. Some species have small, dispersed segments of introgression left over from historical hybridization events (Edelman et al., 2019; Green et al., 2010; Nelson et al., 2021; Randi & Lucchini, 2002). Others form hybrid swarms with a well-mixed continuum of ancestry (Behm et al., 2010; Hasselman et al., 2014; Kleindorfer et al., 2014; Ruhsam et al., 2011). In tension zones, hybrid ancestry diffuses outward from narrow regions of contact into allopatric regions (Barton & Hewitt, 1985, 1989; Macholán et al., 2007; Martinsen et al., 2001; Moore & Buchanan, 1985; Streisfeld & Kohn, 2005). Other systems form mosaic hybrid zones, with variable patches of contact across a heterogeneous landscape (Curry, 2015; Howard et al., 1993; Rand & Harrison, 1989; Valbuena-Carabaña et al., 2007).

Demographic and stochastic forces generate variation in introgression across individuals and the genome. Different population sizes, dispersal rates, and expansions or bottlenecks help determine the relative contribution of two species to admixed genomes (Bourret et al., 2022; Klein et al., 2017; Quilodrán et al., 2020; Secondi et al., 2006; Valbuena-Carabaña et al., 2007). Mating dynamics can lead to asymmetries in the direction of introgression (Kenney & Sweigart, 2016; N. H. Martin et al., 2025; Pickup et al., 2019; Ruhsam et al., 2011; Sianta et al., 2024; Sweigart & Willis, 2003). The time since hybridization, and whether it happens in a single pulse or continuously over time, will affect the size distribution of introgressed ancestry blocks (Menon et al., 2021; Verdu & Rosenberg, 2011).

Beyond these demographic forces, selection can have strong effects on the distribution of introgression (Sachdeva & Barton, 2018; Sedghifar et al., 2016). A common theme is pervasive, polygenic selection in hybrids against ancestry from the minor parent (i.e., the parent contributing the smaller fraction of hybrid genomes) (Juric et al., 2016; Schumer et al., 2018). This could be due to higher genetic load in that species (Juric et al., 2016; Nouhaud et al., 2022), intrinsic incompatibilities between genomes (Schumer et al., 2018), or local adaptation and coadaptation favoring major-parent alleles (Hohenlohe et al., 2012). In some systems, a correlation between introgression and recombination rate has been cited as evidence for pervasive selection against minor-parent ancestry (Brandvain et al., 2014; Burri et al., 2015; S. H. Martin et al., 2019; Nouhaud et al., 2022; Schumer et al., 2018), because selection in lower-recombination regions can efficiently remove larger blocks of linked hybrid ancestry, lowering effective introgression rates. The process is analogous to background selection on deleterious alleles within species, which can lead to a correlation between nucleotide diversity and recombination rate (Charlesworth et al., 1993; Comeron, 2014; Cruickshank & Hahn, 2014; Stephan, 2010). However, many plant species do not show this within-species background selection pattern (Fernandes et al., 2024; Lovell et al., 2025; Slotte, 2014), possibly because other factors, such as an association between gene density and recombination rate, can act to attenuate the signal of linked selection. It remains unresolved whether linked selection against hybridization will follow the same relationships as within-species background selection in such cases, or whether linked selection after hybridization has unique characteristics.

In addition to genome-wide patterns, specific loci may be under selection in admixed populations, leading to detectable signals of either reduced or elevated introgression. Areas of reduced introgression have been associated with genes controlling intrinsic incompatibilities (Moran et al., 2024; Powell et al., 2020; Runemark et al., 2018). However, preferential introgression of compatible alleles across species boundaries can also collapse or eliminate potential incompatibilities, hiding them from detection (Bank et al., 2012; Frayer & Payseur, 2024; Lemmon & Kirkpatrick, 2006; Xiong & Mallet, 2022). Locally adapted alleles are another possible source of loci resistant to introgression (Jones et al., 2012; Lowry & Willis, 2010). Expectations involving premating reproductive barrier loci are less clear: in theory, assortative mating loci should resist introgression (Streisfeld & Kohn, 2005; Wessinger et al., 2023), but mating-related loci are also documented crossing species boundaries and fixing in new lineages (Nelson et al., 2021; Stankowski & Streisfeld, 2015). Meanwhile, areas of elevated minor-parent ancestry might indicate cases of adaptive introgression (Chhatre et al., 2018; Pardo-Diaz et al., 2012; Racimo et al., 2015), in which the minor parent allele is favored either globally or in a specific admixed environment. Another source of elevated minor-parent ancestry could be the introgression of selfish genetic elements, which are expected to easily cross species boundaries when given the opportunity (Rushworth et al., 2022; Sweigart et al., 2019).

When hybridization occurs multiple times across a landscape, comparing and contrasting these natural replicates can highlight the underlying processes involved. A few studies have shown remarkable similarity in independent cases of hybridization of the same species pair (Chaturvedi et al., 2020; Nouhaud et al., 2022). Species pairs with higher divergence prior to hybridization appear to have more repeatable genomic patterns of introgression in secondary contact than more closely related species pairs (Langdon et al., 2024). Variation across space can be driven by demographic factors such as population expansion or invasion (Quilodrán et al., 2020), or gene flow filtering outward from the center of a contact zone (Martinsen et al., 2001; Simon et al., 2021). But the underlying factors influencing predictability and parallelism in hybridization remain underexamined.

We might expect that loci identified by mapping studies as important for reproductive isolating barriers would have predictable responses in ancestry frequencies within wild populations. However, there are few direct tests of the correspondence between these two types of data, in part because few systems have both types of data available. Where differentiation in hybrid zones has been associated with known adaptive or barrier loci, it is usually in the context of chromosomal inversions that suppress recombination (Dean et al., 2024; Machado et al., 2007) or genes on sex chromosomes (Payseur & Nachman, 2005; Putnam et al., 2007; Trier et al., 2014). A recent analysis of the *Mus musculus musculus x M. musculus domesticus* hybrid zone and corresponding laboratory crosses found that wild and laboratory studies identified distinct and rarely overlapping sets of loci (Frayer & Payseur, 2024). A multispecies analysis in cichlids also found low correspondence between different methods of detecting putative incompatibilities (Feller et al., 2024).

The *Mimulus guttatus x M. nasutus* study system is ideal for addressing broad questions about the nature of hybridization and introgression. Hybridization has been detected in multiple distinct geographic areas (Brandvain et al., 2014; Kenney & Sweigart, 2016; Zuellig & Sweigart, 2018a), allowing for comparative tests of genomic introgression patterns across these replicates. Furthermore, multiple premating and postmating reproductive incompatibilities have been mapped to candidate loci (Fishman et al., 2014; Zuellig & Sweigart, 2018b) or quantitative trait loci (Mantel & Sweigart, 2024), setting us up to test the correspondence between introgression patterns in hybrid populations and mapped reproductive isolation traits.

In this study, we take advantage of previously generated sequencing data from two hybridizing populations in Washington, USA: Catherine Creek (CAC) and Little Maui (LM) (Farnitano et al., 2025). These populations each consist of an asymmetric and dynamic swarm of hybrids with *M. guttatus* as the major and *M. nasutus* as the minor parent, shaped by partial connectivity with co-occuring parental species (Farnitano et al., 2025; Kenney & Sweigart, 2016). We combine those data with new data from a second, completely independent region of hybridization, the foothills and montane regions of north-central California, USA. By comparing patterns of genomic ancestry within and among these regions, we seek to illuminate how diverse processes shape introgression. We focus on four key questions: A) How similar are hybrid ancestry outcomes and genomic patterns of ancestry across different hybridizing populations? B) To what extent are shared patterns of ancestry driven by features of the genomic landscape versus idiosyncratic patterns at individual loci? (C) How much does migration between nearby populations influence the distribution of introgression? (D) Do candidate loci known to influence reproductive isolation show predictable and parallel patterns of genomic ancestry? Each of these questions is key to our understanding of the selective processes occurring during and after hybridization, with implications for predicting responses to hybridization across the tree of life.

## MATERIALS AND METHODS

### Study sites and sample collection

We collected seeds or leaf tissue from individuals from 21 locations across the foothills and mountains of the Sierra Nevada in northern California, USA (Southern region, Figure 1A,C). Southern region samples were collected in the field from April to June of 2021. We visited most locations multiple times to capture a range of phenological variation, targeting our sampling to obtain both *M. nasutus* and *M. guttatus* as well as any likely hybrids or admixed individuals, including sampling both senesced and actively flowering individuals where available. However, we note that sampling was not exhaustive and may not be numerically representative of the proportions of *M. guttatus* vs. *M. nasutus* at any one location.

**Figure 1.**
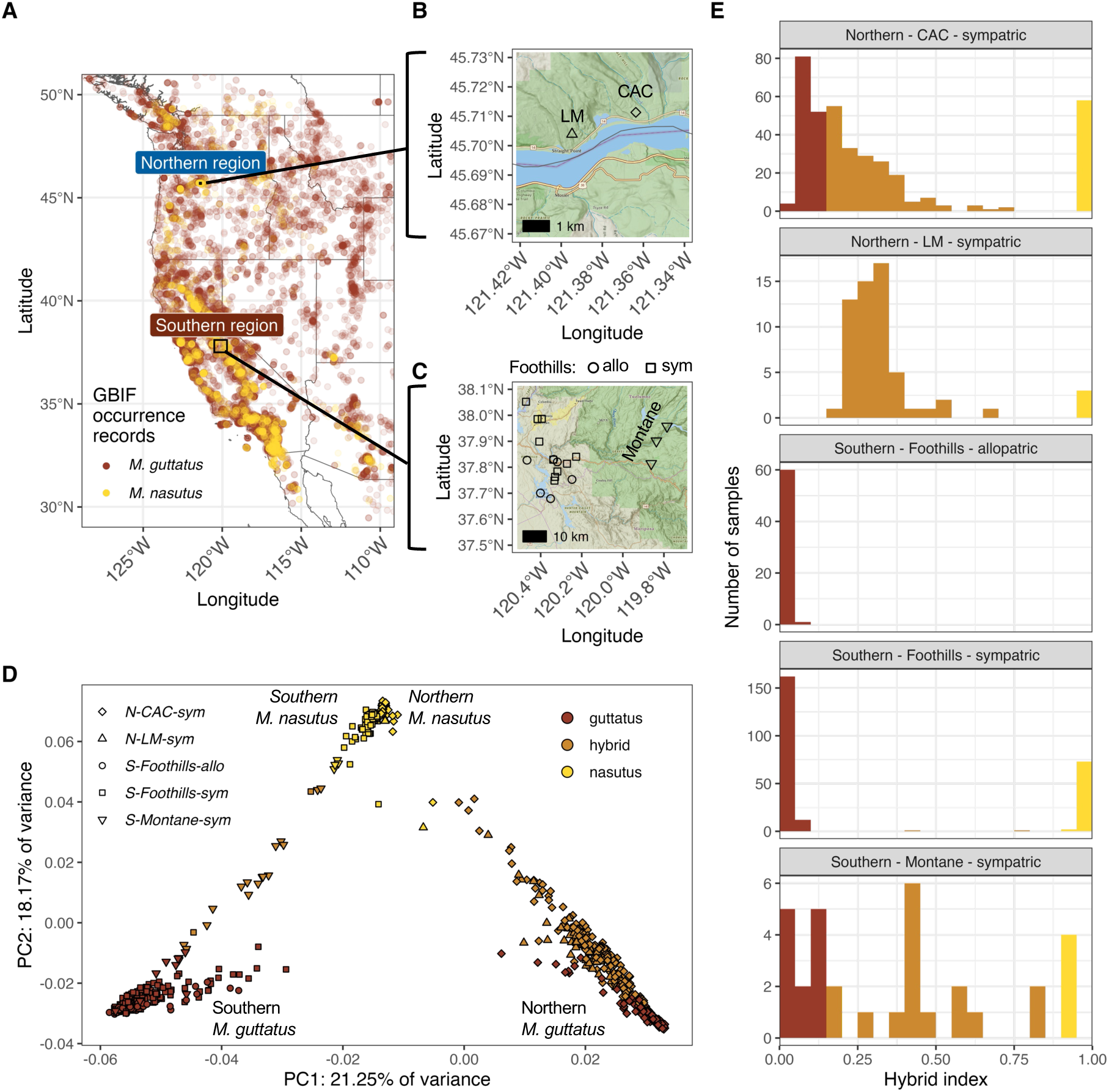
Range map and population structure of sampled contact populations between *M. guttatus and M. nasutus*. (A) Range map of *Mimulus guttatus* (red) and *Mimulus nasutus* (yellow) observations in the western United States, highlighting focal Northern and Southern sampling regions for this study. Points are occurrence records from GBIF(GBIF.org User, 2025). (B) Location of two sympatric sampling locations in the Northern region, Catherine Creek (CAC) and Little Maui (LM), along the Columbia River in southern Washington, USA. (C) Location of sampling locations in the Southern region, including five allopatric *M. guttatus* Foothills populations, ten Foothills sympatric populations where both *M. nasutus* and *M. guttatus* were collected, and three Montane (>1000m) sympatric populations in the Sierra Nevada mountains. Map images in B and C were obtained from the NAIP public image services(U. S. Department of Agriculture, Farm Service Agency (FSA) & Aerial Photography Field Office (APFO), 2025). (D) Genomic principal components analysis of genotype likelihoods for samples in this study, using the PCAngsd method. Regional differentiation occurs along PC axis 1, while interspecific differentiation appears along PC axis 2. (E) Histogram of hybrid index, the proportion of genomic ancestry assigned to *M. nasutus* for each individual, among samples from each of five geographic groupings. Hybrid index of 0.0 indicates 100% *M. guttatus* ancestry, while hybrid index of 1.0 indicates 100% *M. nasutus* ancestry. Ancestry assignments are from ancestryHMM outputs binned into 50kb genomic windows. Colors indicate the three cohort designations used in downstream analysis: *guttatus* (HI<0.15), *hybrid* (HI 0.15-0.85), and *nasutus* (HI>0.85).

Tissue from wild-growing individuals was dried with silica gel beads in the field. For wild-collected seeds, a subset of seeds from each maternal family were germinated in the UGA Botany greenhouses, where bud or leaf tissue was collected from a single germinant per maternal family and stored at -80C until the time of DNA extraction. For all samples, DNA was flash-frozen in liquid nitrogen, ground with metal beads, and extracted using a CTAB protocol with phenol-chloroform extraction (Fishman, 2020). Samples were prepped for low-coverage whole-genome short-read sequencing using a custom tagmentation library prep (Farnitano & Sweigart, 2024) with bulk-prepped Tn5 tagmentation enzyme (Lu et al., 2017), following the same procedure used for earlier CAC and LM samples (Farnitano et al., 2025). Samples were sequenced on an Illumina NovaSeq 6000 machine at the Duke University Sequencing and Genomic Technologies Core Facility with 150bp paired reads to an average depth of ∼1.25 million paired reads per sample (median 0.99 million read pairs). We sequenced a total of 382 individuals from the Southern region (Table S1).

We combined these Southern region samples with 503 previously sequenced (Farnitano et al., 2025) individuals from two Northern sites in Washington, USA: Catherine Creek (CAC), and Little Maui (LM), consisting of *M. nasutus, M. guttatus*, and admixed individuals (Table S1). CAC individuals were collected as wild tissue samples from across the growing season over three years: 2019, 2021, and 2022; LM individuals were collected only in 2021.

### Ancestry reference panel and SNP marker lists

To obtain a set of ancestry-informative markers for distinguishing *M. guttatus* and *M. nasutus* ancestry, we genotyped a panel of 33 geographically diverse inbred lines: 5 *M. nasutus*, 9 *M. guttatus* from the ‘Northern’ clade, 11 *M. guttatus* from the ‘Southern’ clade, and 8 admixed *M. guttatus* from the CAC population (Table S2; See Supplementary Text for genotyping details). As a high-quality marker list, we chose biallelic SNPs with a called genotype in >=80% of samples from each subgroup within the panel (i.e., *M. nasutus,* Northern *M. guttatus*, Southern *M. guttatus,* and CAC admixed *M. guttatus*).

For population structure inference, we randomly thinned this marker list to one SNP per 1000bp, resulting in 79,966 *angsd* markers. For ancestry inference, we did not thin, but instead selected SNPs with an allele frequency difference between *M. guttatus* panel members and *M. nasutus* panel members of at least 80% (excluding the admixed CAC individuals), resulting in 201,387 ancestry-informative markers.

### Genomic PCA analysis

To assess population structure and variation across northern and southern samples, we used *angsd v0.940* (Korneliussen et al., 2014) with the GATK likelihood method to estimate genotype likelihoods at each *angsd* marker for each individual. Low-coverage samples may be sensitive to false homozygosity if overlapping forward and reverse reads are counted twice during genotype likelihood estimation, so we first merged overlapping forward and reverse reads using *Flash v 2.2.00* (Magoč & Salzberg, 2011). Then, we trimmed, mapped, and filtered reads using the same approach and parameters as the reference panel (See Supplementary Text), and used the resulting alignments as input for *angsd* genotype likelihood estimation. We used the genotype likelihoods to conduct a principal components analysis (PCA) in *PCAngsd v1.10* (Meisner & Albrechtsen, 2018). For the final dataset, we excluded one individual from the Southern dataset that was discovered to be a contaminant, as it clustered with Northern individuals in our initial PCA.

### Local ancestry inference

We followed the approach of *ancestryinfer* (Schumer et al., 2020) using *Ancestry_HMM* (R. Corbett-Detig & Nielsen, 2017) to estimate local ancestry (*M. guttatus, M. nasutus,* or heterozygous) across the genome for each reference panel member or low-coverage Northern and Southern sample. Briefly, samples were aligned to the *M. guttatus* IM767v2 reference genome and the *M. nasutus* SFv2 reference genome (Lovell et al., 2025), keeping only reads that aligned exactly once to each genome. We then used the alignments to the IM767v2 reference to extract read counts per sample at each ancestry-informative marker (defined above); note that each read pair is only counted once here. Posterior probabilities for ancestry assignment were obtained with *Ancestry_HMM,* and ancestry genotypes were called for all markers with a posterior probability >=0.95. We then used a windowing approach to smooth and filter ancestry calls. For each 50kb non-overlapping genomic window, we assigned an ancestry genotype to the window if at least five sites within the window had called ancestry genotypes AND if >=90% of those called sites agreed on their ancestry call; the remaining window ancestry genotypes were set to missing.

We removed from the dataset any windows where fewer than 50% of samples had a called ancestry genotype, then removed any sample with genotype calls in fewer than 50% of the remaining windows. Our final filtered dataset included 781 individuals (437 Northern and 344 Southern) with ancestry genotypes at 3,140 genomic windows. For each individual, we calculated a hybrid index: the genome-wide proportion of *M. nasutus* ancestry alleles within called window genotypes. We classified each individual into three cohorts according to their hybrid index: HI < 0.15 = *guttatus*, HI 0.15-0.85 = *hybrid*, and HI > 0.85 = *nasutus*.

### Ancestry frequencies and their correlations across groups

To facilitate comparisons among samples, we divided our dataset into five geographic groupings: the Northern CAC sympatric (N-CAC-sym) population; the Northern LM sympatric (N-LM-sym) population; a Southern montane sympatric (S-Montane-sym) group consisting of samples from three sympatric locations above 1000m elevation in the Sierra Nevada mountains; a Southern foothills sympatric (S-Foothills-sym) group consisting of Southern locations below 1000m elevation where both *M. guttatus* and *M. nasutus* were sampled; and a Southern foothills allopatric (S-Foothills-allo) group consisting of all Southern locations where only *M. guttatus* individuals were found. Note that we sampled no locations with only *M. nasutus* samples, and no allopatric Montane populations. Within each geographic grouping, we divided samples further into three ancestry cohorts as above, resulting in 15 potential geographic-by-ancestry sample groups. Some groupings did not have sufficient representation from all cohorts; we focused our comparative analyses on nine geographic-by-ancestry groups with at least 10 samples (Table S1). For each of these nine groups, we calculated the frequency of *M. nasutus* alleles in the group at each 50kb genomic window (ancestry frequencies).

To address the overall level of parallelism in ancestry, we calculated the proportion of shared variance (*R^2^*) in ancestry frequencies across all 50-kb genomic windows for each pair of geographic-by-ancestry groups.

### Effect of genomic features and missing data on ancestry frequencies

We obtained values for chromosomal position, gene density, and recombination rate to investigate their relationships with ancestry frequencies across the genome. We calculated a ‘squared relative chromosomal position’ – the square of the distance in base pairs from the center of each 50kb genomic window to the midpoint of the chromosome reference sequence, scaled by the size of each chromosome (squared distance provided a better fit to ancestry proportions than absolute distance). We obtained the number of annotated genes falling within each 50kb genomic window of the IM767 v2 reference (Lovell et al., 2025). We obtained recombination rates by taking a dataset of 33302 *M. guttatus* recombination events from an F2 panel (Veltsos & Kelly, 2024) and assigning each event to a 50kb genomic window. Because recombination rates were locally noisy at this 50kb resolution, we also calculated smoothed rates by grouping each window into a nonoverlapping 1Mb window and taking the average recombination rate. To obtain a proxy for the strength of linked selection, we divided the number of genes per window (i.e., the density of targets of selection) by the recombination rate (using the 50kb-resolution value).

In addition to investigating the influence of genomic features, we explored whether differences in data quality across the genome affected our assessments of hybrid ancestry. Mapping bias or other technical artifacts could cause certain genomic windows to be biased towards one genotype call over the other; these windows might also have lower genotype calling confidence and therefore higher numbers of samples with missing data. To obtain a proxy for data quality and the reliability of ancestry calling, we calculated the fraction of samples with missing data at each genomic window. We calculated this ‘missingness’ fraction separately for each sample group; note that windows with missingness greater than 0.5 were already removed entirely, as described above.

For each of the genomic features described above, including missingness, we calculated its individual correlation with the ancestry frequencies of each sample group. Then, to assess the combined effect of these features, we ran a type-I analysis of variance (ANOVA) for each sample group. We used a linear model with ancestry frequency as the dependent variable, and the following independent variables: squared relative chromosomal position, gene density, recombination rate (at 1Mb resolution), recombination rate (at 50kb resolution), and missingness proportion. The residuals from this model are used in further analyses described below, and referred to as ‘residual ancestry frequencies.’

Next, for each pair of sample groups, we assessed how much of the remaining variance in ancestry frequency for one sample group is still explained by the ancestry frequency of the other group after accounting for genomic features and missingness. We ran a corresponding type-I ANOVA, using the same linear model above with ancestry frequency of the first group as the dependent variable and genomic features and missingness as independent variables, but with an additional independent variable term for the ancestry frequency of the second group. Note that no more than two sample groups (one dependent and one independent) were included in any given model.

### Ancestry frequency outliers as a test of shared positive selection

We used outlier windows with high minor-parent ancestry frequency to represent preferentially introgressing loci, in order to assess whether positive selection significantly impacts shared patterns of ancestry across groups. For each sample group, we designated the 50kb windows in the top 5% of minor-parent ancestry frequency (high *M. nasutus* ancestry frequency in *guttatus* or *hybrid* groups, and low *M. nasutus* ancestry in *nasutus* groups) as ‘uncorrected’ outlier windows. Similarly, we identified outliers for ‘residual ancestry frequencies’ from the linear model described above, which are corrected for genomic features and missingness, designating the top 5% of minor-parent residual ancestry frequencies as ‘model-corrected’ outlier windows. For both the uncorrected and model-corrected outliers, we counted the number of overlapping outlier windows between each pair of sample groups. To test significance of the overlap, we used a chi-squared test for a deviation from the expected overlap if windows were picked randomly, applying a Bonferroni correction to account for multiple tests.

### Ancestry frequency effects at known reproductive isolation candidate loci

We used existing candidate loci to test whether reproductive isolation loci from laboratory crosses have predictable and parallel effects on ancestry frequencies. We first used a dataset of 20 quantitative trait locus confidence intervals representing putative intrinsic reproductive-isolation-associated loci, derived from three recombinant-inbred-line mapping populations between *M. guttatus* and *M. nasutus* lines from the Northern CAC population (Mantel & Sweigart, 2024). These included twelve putative incompatibility regions associated with conditional reductions in male or female fertility, plus eight regions associated with epistatic transmission ratio distortion; two additional pairs of transmission-ratio-distortion regions from the original dataset were excluded because their confidence intervals exceeded half of a chromosome. For the boundaries of each interval, we used a 1.5-LOD drop from the peak LOD score. We designated each locus as ‘N-’ or ‘N+’ based on the predicted direction of effect (Mantel & Sweigart, 2024): for the seven N- loci, the *M. nasutus* allele is associated with relatively lower fertility or is under-transmitted in mapping lines, so we hypothesize an decrease in wild *M. nasutus* frequencies; while for the thirteen N+ loci, the *M. guttatus* allele is associated with relatively lower fertility or is under-transmitted in mapping lines, so we hypothesize an increase in wild *M. nasutus* frequencies.

We tested whether 50kb windows falling within N- locus (or N+ locus) intervals had significantly lower (or higher) residual ancestry frequencies compared to the genome-wide average, using individual one-tailed t-tests for each sample group, with a Bonferroni correction to account for multiple tests.

Next, we tested whether our uncorrected or model-corrected ancestry outlier windows fell within these N- or N+ QTL intervals more often than expected by chance. We counted the number of outliers in QTL intervals, then used a chi-squared test to assess the deviation from a null expectation of outliers in QTL intervals if the outlier windows were chosen randomly.

Finally, we assessed the behavior of ancestry frequencies around four additional reproductive isolation loci that have been fine-mapped to individual candidate genes or gene clusters: two classical Dobhzansky-Muller incompatibility loci that interact to produce hybrid lethality (Zuellig & Sweigart, 2018a, 2018b), and two major QTL impacting photoperiod between *M. guttatus* and *M. nasutus* (Fishman et al., 2014). We assigned each candidate to a single 50kb window in the IM767v2 reference assembly, then compared ancestry at these windows to the genome-wide distribution within each sample group, asking whether they coincided with relative peaks or troughs of ancestry. For each focal window, we calculated its quantile within the genome-wide distribution of both raw and residual (model-corrected) ancestry frequencies, as well as the average quantile of the surrounding neighborhood, defined as the focal window plus all windows within 500kb on either side.

We used the *stats* and *tidyverse* packages in R (R Core Team, 2023; Wickham & RStudio, 2023) for all statistical inference and data visualization.

## RESULTS

### Multiple regions of contact between species have resulted in a wide variety of hybridization outcomes

In total, we obtained ancestry genotype assignments for 782 individuals across 3,140 50kb genomic windows. Genomic PCA analysis indicated both geographic and species-level clustering: PC axis 1 (21% of variance) separated Northern and Southern *M. guttatus,* with a more subtle but parallel separation of Northern and Southern *M. nasutus*; PC axis 2 (18% of variance) separated *M. guttatus* from *M. nasutus,* with a large number of intermediate hybrid individuals in a continuum between species for both the Northern and Southern groups (Figure 1D).

We found detectable variation in hybrid ancestry within all five geographic groupings, even the allopatric Southern foothills group (Figure 1E). Consistent with previous findings, the Northern populations were asymmetric, with robust *hybrid* cohorts skewed towards majority-*M. guttatus* ancestry. All Northern *guttatus* individuals retained at least some *M. nasutus* ancestry (HI>0.01) indicating pervasive directional introgression. In contrast, we found only a few *hybrid* cohort individuals within Southern Foothills sympatric populations. Still, *guttatus* HI values were frequently greater than 0.01, indicating small but detectable introgression of *M. nasutus* ancestry (137 of 147 individuals). In the Southern Foothills allopatric populations, HI values were typically less than 0.01, but we found a few individuals (12 of 60) with HI>0.01, up to a maximum of HI=0.08, demonstrating residual *M. nasutus* introgression in these populations. Finally, in the Southern montane sympatric group we found a wide range of HI values with a comparatively symmetric distribution, including some *M. nasutus* with small amounts of *M. guttatus* ancestry (four individuals with HI 0.9-0.95).

### Ancestry frequencies are highly variable across the genome and strongly correlated among hybridizing populations from distant regions

Ancestry frequencies varied substantially across the genome in all groups (Figure 2A). For *hybrid* cohorts, ancestry frequencies covered the full range from near 0.0 to near 1.0, but were generally asymmetric in the North, with more windows in lower (majority-*guttatus*) frequencies. Southern montane *hybrids* were less asymmetric; a large number of windows at 50% frequency suggests the presence of early-generation hybrids, contrasting with the later-generation hybrids indicated by a wider distribution of frequencies in the North. For all *guttatus* groups, even the Southern Foothills allopatric group, there were genomic windows with substantial *M. nasutus* ancestry, though the extent varied across groups. The Northern CAC *guttatus* group, strikingly, had windows with up to 88% *M. nasutus* ancestry; the Southern Foothills sympatric *guttatus* group had windows with *M. nasutus* ancestry as high as 35%. Meanwhile, both Northern and Southern *nasutus* cohorts had very few windows with any detectable *M. guttatus* ancestry, consistent with strongly asymmetric introgression. However, there were a few windows with evidence of *M. guttatus* introgression into *M. nasutus*: ancestry frequencies reached as low as 0.15 (i.e., 85% *M. guttatus*) and 0.52 (48% *M. guttatus*) for Northern CAC *nasutus* and Southern Foothills *nasutus,* respectively.

**Figure 2.**
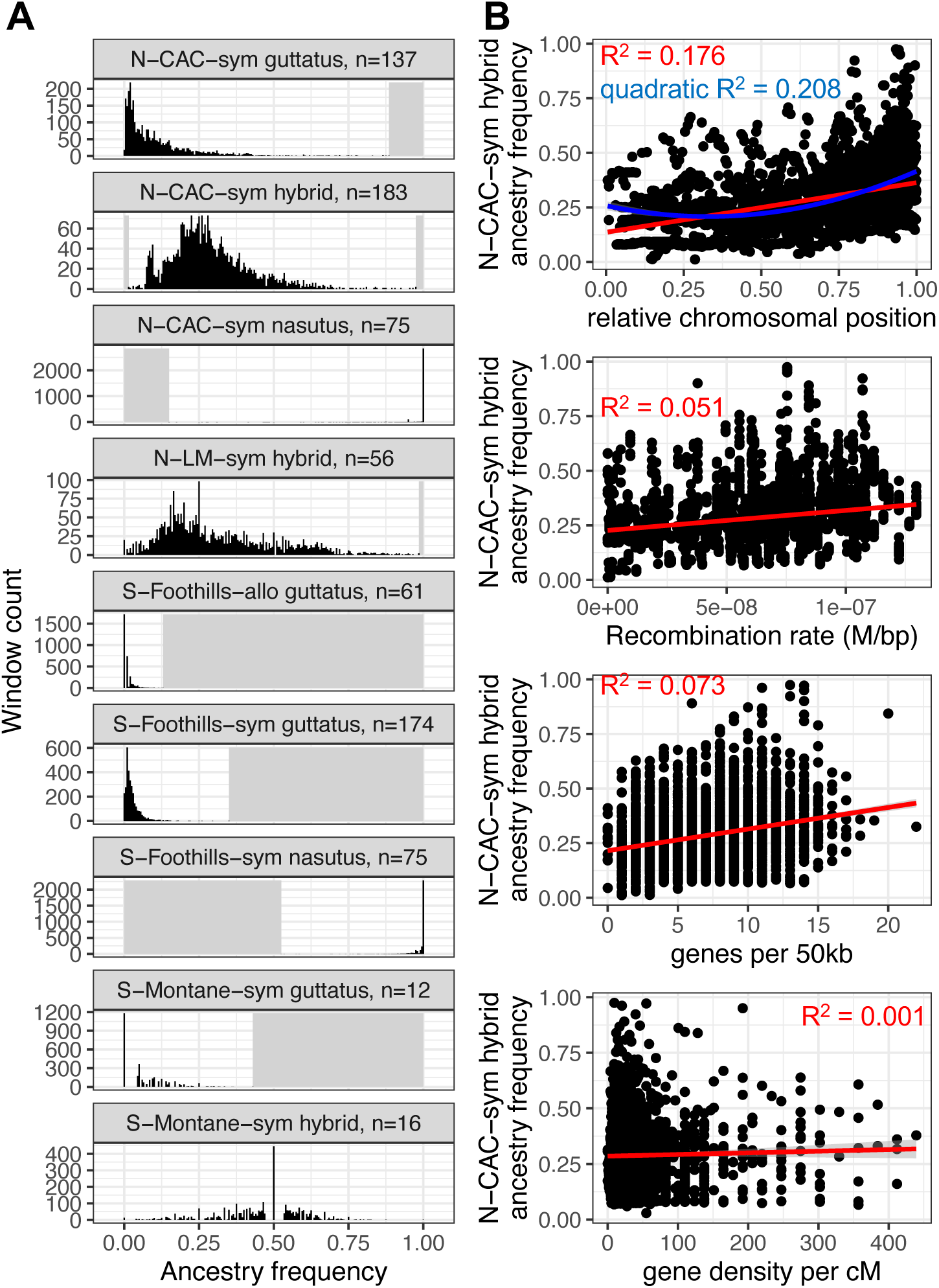
Wide distribution of ancestry frequencies across 50kb genomic windows and their correlation with genomic features. (A) Histogram of ancestry frequencies across 3,140 50kb genomic windows for each of eight geographic x ancestry sample groups. N-CAC-sym = Northern Catherine Creek sympatric, N-LM-sym – Northern Little Maui sympatric, S-Foothills-allo = Southern Foothills allopatric, S-Foothills-sym = Southern Foothills sympatric, S-Montane-sym = Southern Montane sympatric; individuals with hybrid index 0-0.15 are categorized as *guttatus,* HI 0.15-0.85 as *hybrid,* and HI 0.85-1.0 as *nasutus*. Ancestry frequency is the proportion of *M. nasutus* ancestry among called samples for a given window. Values below the minimum or above the maximum frequency for each group are shaded with grey. (B) Scatter plots indicating correlations between genomic features and ancestry frequencies of 50kb windows for one representative sample group, N-CAC-sym *hybrid*. Relative chromosomal position is the position of each window within its chromosome, ranging from 0.0 at the midpoint to 1.0 at either end. Recombination rate is shown here at 1Mb resolution; R^2^=0.013 for recombination rate at 50kb resolution in this sample group. Genes per 50kb is based on counts of annotated genes within each 50kb window; genes per cM is the same value scaled by the recombination rate at 50kb resolution. Red lines and adjusted-R^2^ values are from a simple linear model y∼x; blue line and quadratic adjusted-R^2^ for the top panel is from a simple linear model y∼(x^2).

For comparisons between any two *guttatus* or *hybrid* groups, the ancestry frequencies of one group explained a substantial amount of the genomic variation in ancestry frequencies of the other group, indicating substantial genomic parallelism (Figure 3, bottom right triangle). This effect was strongest within populations or between populations in the same geographic region, but even extended to comparisons between Northern and Southern regions. In contrast, ancestry frequencies in *nasutus* cohorts were typically not strongly correlated with *guttatus* or *hybrid* cohorts, but were correlated with each other between North and South.

**Figure 3.**
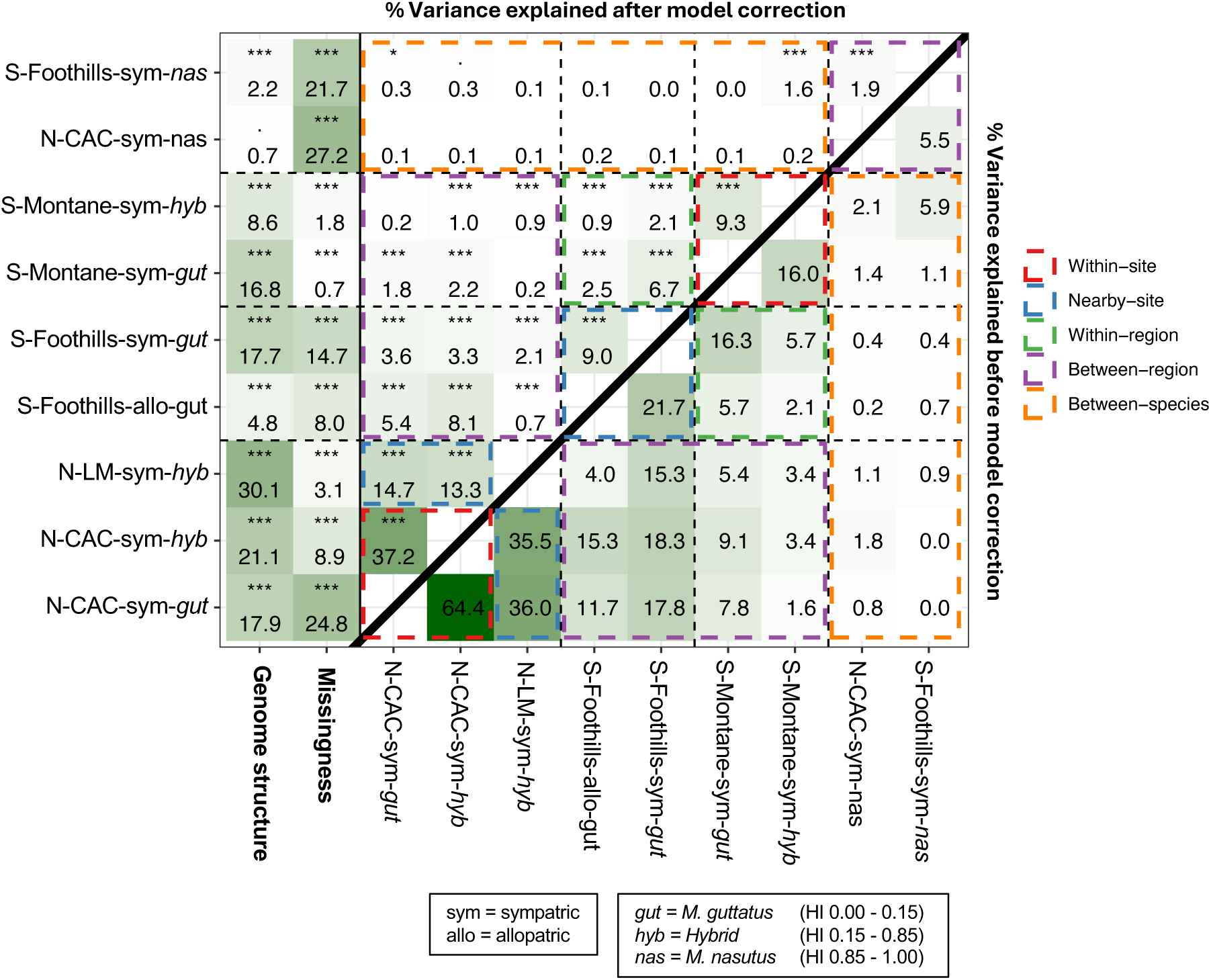
Substantial correlations among ancestry frequencies across geographic regions are explained in part by genomic and technical features. Bottom right triangle: percent of variance in ancestry frequency among windows in sampling group Y (y-axis) explained by ancestry frequency in sampling group X (x-axis), under a simple linear model Y ∼ X with no corrections for other factors. In this simple case, variance is equal to the correlation coefficient R^2^ between Y and X. Note that each pairwise comparison is calculated independently; rows or columns are not additive. Top left triangle: percent variance in ancestry frequency in sampling group Y (y-axis) explained sequentially (using a type-I ANOVA) by genome structure, missingness, and sampling group X (x-axis), under a full linear model Y ∼ [genomic features] + missingness + X. Note that separate models are run for each pairwise sampling group comparison, so variance attributed to each group is not additive. Genomic features included the squared distance to the chromosome midpoint, gene density, and recombination rate at both 1Mb and 50kb resolutions; variance components for these features were added together. Missingness refers to the within-group fraction of individuals with a missing ancestry genotype at each window. Comparisons are grouped by colored rectangles according to their comparison type: within collection locations, between nearby locations, between more distant groups within the same region, and between Northern and Southern regions; comparisons involving *nasutus* cohorts are grouped separately.

### Genomic structural features and technical factors, but not linked selection, explain a large proportion of the shared variance in ancestry frequencies across groups

The squared distance to the chromosome center, the density of genes within a genomic window, and the recombination rate (calculated at two resolutions, 1Mb and 50kb) were all significantly correlated with ancestry frequencies in most groups (Figure 2B, Table 1, Table S3). These features were all also correlated to each other, making their individual effects difficult to separate. However, chromosomal position and gene density were in most cases better predictors of ancestry than recombination rate (Figure 2B, Table 1, Table S3). These measures also retained their relationship if centromeric regions were excluded, whereas the association with recombination rate was substantially weaker outside centromeric regions (Figure 2B, Table S3). Surprisingly, the density of genes scaled by the recombination rate, a more direct proxy for the effect of linked selection, was only weakly if at all correlated with ancestry frequency (max R^2^=0.008, or 0.018 excluding pericentromeric regions, Figure 2B, Table 1, Table S3). These findings suggest that, while structural features as a whole are predictive of hybrid ancestry, their association is not a direct result of linked selection as argued in other systems (Burri et al., 2015; S. H. Martin et al., 2019; Nouhaud et al., 2022; Schumer et al., 2018).

**Table 1.** Correlation coefficients (r) between genomic features and ancestry frequencies in 50kb windows.

|  | Relative<br>position | Relative<br>position<sup>2</sup> | Gene<br>density<br>(per bp) | Recomb.<br>rate (1Mb) | Recomb.<br>rate (50kb) | Gene<br>density<br>(per cM) |
| --- | --- | --- | --- | --- | --- | --- |
| Relative position | 1 | 0.977 | 0.540 | 0.609 | 0.329 | -0.245 |
| Relative position <sup>2</sup> | 0.977 | 1 | 0.554 | 0.560 | 0.298 | -0.199 |
| Gene density (per bp) | 0.540 | 0.554 | 1 | 0.575 | 0.329 | -0.101 |
| Recomb. rate (1Mb) | 0.609 | 0.560 | 0.575 | 1 | 0.548 | -0.326 |
| Recomb. rate (50kb) | 0.329 | 0.298 | 0.329 | 0.548 | 1 | -0.170 |
| Gene density (per cM) | -0.245 | -0.199 | -0.101 | -0.326 | -0.170 | 1 |
| *Ancestry frequency* |  |  |  |  |  |  |
| N-CAC-sym *guttatus* | 0.384 | 0.416 | 0.272 | 0.204 | 0.114 | -0.053 |
| N-CAC-sym *hybrid* | 0.420 | 0.457 | 0.271 | 0.227 | 0.115 | -0.068 |
| N-CAC-sym *nasutus* | 0.045 | 0.050 | 0.060 | 0.079 | 0.059 | -0.045 |
| N-LM-sym *hybrid* | 0.498 | 0.543 | 0.337 | 0.272 | 0.122 | -0.083 |
| S-Foothills-allo *guttatus* | 0.171 | 0.218 | 0.120 | 0.116 | 0.049 | -0.003 |
| S-Foothills-sym *guttatus* | 0.376 | 0.408 | 0.309 | 0.268 | 0.138 | -0.069 |
| S-Foothills-sym *nasutus* | 0.106 | 0.104 | 0.117 | 0.140 | 0.098 | -0.052 |
| S-Montane-sym *guttatus* | 0.357 | 0.358 | 0.315 | 0.346 | 0.232 | -0.108 |
| S-Montane-sym *hybrid* | 0.192 | 0.211 | 0.219 | 0.272 | 0.198 | -0.075 |

\*Relative position = (2 \* bp distance from chromosome center) / (chromosome length)

The proportion of missing data in a window was also correlated with ancestry frequency for most groups: when included as a model factor, missingness explained between 0.7% and 27.2% of variation in ancestry frequency, with a wide range across groups (Figure 3, left columns). This suggests that technical variation in ancestry assignment may have an impact on ancestry frequencies, through window-specific biases in ancestry assignment. However, this signal could also represent a real biological signal, in which some unidentified genomic process produces both lower-quality ancestry calls and higher rates of introgression. As an additional test for the effect of technical variation in ancestry calling, we removed windows with low correspondence between ancestry assignment and local genomic PCA structure (Supplementary Text, Figures S1-S6); this analysis resulted in slightly lower correlations between groups overall, but significant variance in ancestry was still explained by ancestry patterns of other groups, particularly for groups in close geographic proximity.

### Geographic proximity is a major source of parallel patterns in hybrid ancestry

After applying a correction for genomic structural features and missingness, we assessed how much of the remaining variance was explained by pairwise comparisons among sample groups (Figure 3, top left triangle). Explained variance was substantially reduced compared to the uncorrected (bottom right triangle) values, but remained significant in most cases. In particular, comparisons between groups connected by gene flow retain the strongest residual correlations: either between cohorts within populations (Figure 3, red rectangles, 8-37% variance explained), or between nearby populations (Figure 3, blue rectangles, 9-14% variance explained). While allele sharing between cohorts within a single population is expected and matches previous findings (Farnitano et al., 2025), correlations between nearby populations appear to indicate a substantial effect of interpopulation gene flow. An interesting case involves the Southern allopatric *M. guttatus* populations, whose ancestry frequencies are best explained by Southern sympatric *M. guttatus* populations (9% of variation explained), suggesting that gene flow from sympatry into allopatry may be spreading introgressed *M. nasutus* alleles. Within allopatry, hybrid ancestry was significantly higher in populations closer to the nearest sympatric location (adj.R^2^=0.73, p=0.009), and was concentrated primarily in three out of six allopatric locations (Figure S7, Table S4). However, correlations between ancestry in allopatry and sympatry persist even when excluding those three locations, showing that introgression extends into all of our sampled populations (Supplementary Text, Figures S8-S9).

### Patterns of shared variation suggest positive selection helps to shape introgression

Between 2.1-3.6% of variation in Southern sympatric *M. guttatus* is explained by ancestry frequencies of Northern groups after accounting for genomic structural features and missingness, despite the ∼1000km distance separating these populations. This residual correlation suggests additional parallel processes shaping ancestry in both groups beyond the structural similarity of their genomes, such as parallel selection for or against particular introgressing loci.

If parallel selection favoring the introgression of particular loci were responsible for some of the correlation in ancestry frequencies among populations, then we hypothesized that outlier windows with unusually high minor-parent ancestry would be shared across populations more often than by chance. To test this hypothesis, we selected the top 5% of windows with the highest minor-parent ancestry for each group (high *M. nasutus* ancestry frequency in *guttatus* or *hybrid* cohorts, and low *M. nasutus* ancestry in *nasutus* cohorts). These outliers overlapped more often than expected by chance in almost all comparisons, with exceptions involving the montane or *nasutus* groups (Figure 4A, bottom right triangle values). Similar to our ANOVA results, the strongest overlaps were seen within geographic groupings or between nearby populations, but some overlap was still found even across Northern vs. Southern regions.

**Figure 4.**
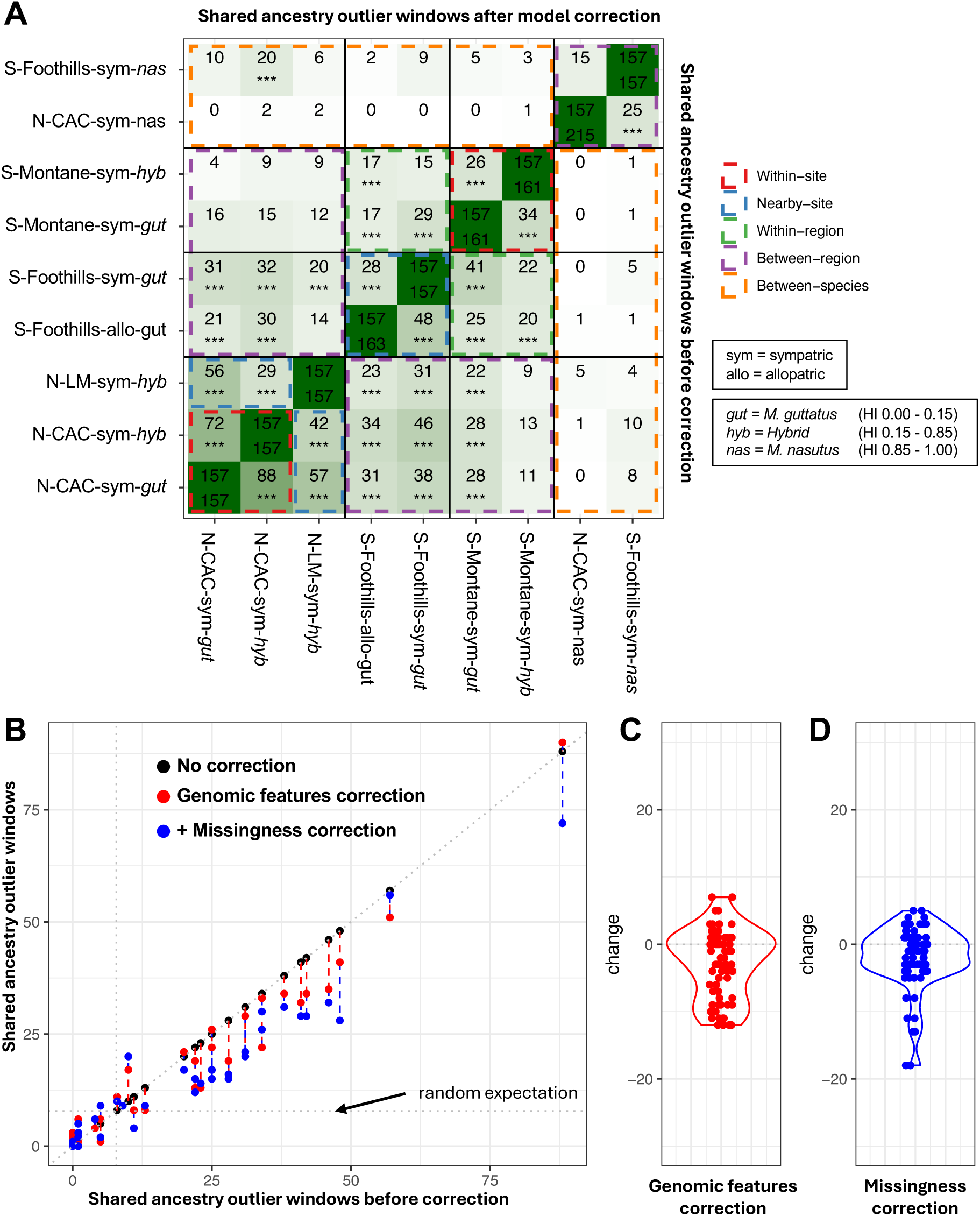
Significant overlap in top-5% ancestry outliers among sampling groups persists after correction for genomic and technical factors. (A) Number of top-5% outlier genomic windows for ancestry that are shared between each pair of sampling groups, using ancestry frequencies prior to any correction (bottom right triangle values) and using ancestry frequency residuals after correction for both genomic features and missingness (top left triangle values). Top-5% outliers were defined as the windows with ancestry frequency equal to or greater than the 95^th^ percentile of ancestry frequency within a sampling group, for *M. guttatus* and *hybrid* groups, and windows equal to or less than the 5^th^ percentile of ancestry frequency, for *M. nasutus* groups. The diagonal indicates the total number of top-5% outliers in each group; note that for uncorrected frequencies, some groups had more than 5%=157 outliers because multiple windows were tied at the 95^th^ or 5^th^ percentile value. A significant deviation from the random expectation (5% of 5%, or 7.85 shared outliers) is indicated with asterisks (***p<0.001 after Bonferroni correction for 72 tests). Comparisons are grouped by colored rectangles as in Figure 3. (B) Change in the number of overlapping outlier windows between groups, after vs. before correction for genomic structural features (the squared distance to the chromosome midpoint, gene density, and recombination rate at both 1Mb and 50kb resolutions), as well as with an additional correction for proportion of samples with missing data. Black dots indicate outlier sharing before correction, red dots indicate sharing after genomic features correction, and blue dots indicate sharing after both genomic features and missingness correction. A dotted line shows the expected overlap if outlier windows were randomly selected. (C and D) Violin plots indicate the difference between the amount of overlap before and after each correction.

To see whether these overlaps are explained predominantly by genomic structural features or missing data, we recalculated outlier windows using residual ancestry frequencies after including these additional factors in a linear model. The degree of overlap was reduced by these corrections, but not eliminated in most cases (Figure 4B, top left triangle values, and Figure C-D). For example, Southern sympatric *M. guttatus* outliers still overlapped significantly with Northern groups.

### Quantitative trait loci and candidate genes associated with reproductive isolation are sometimes, but inconsistently, associated with deviations in ancestry frequency

We next asked whether known reproductive isolation candidate loci were reliable predictors of deviations in ancestry frequency. We first tested 20 quantitative trait loci (QTL) associated with reproductive isolation between *M. guttatus* and *M. nasutus*. For QTL with putatively detrimental *M. nasutus* alleles (N- loci), ancestry frequencies were significantly below predicted values in both Northern population hybrid groups (CAC and LM *hybrids*), but no significant deviation was found for other sample groups (Figure 5, left). For QTL with putatively detrimental *M. guttatus* alleles (N+ loci), ancestry frequencies were significantly above predicted values in some Southern groups (Foothills allopatric and sympatric *M. guttatus*, and Montane *hybrids*) and for CAC *nasutus*, but no significant deviation was found for other sample groups including for CAC and LM *hybrids* (Figure 4, right).

**Figure 5.**
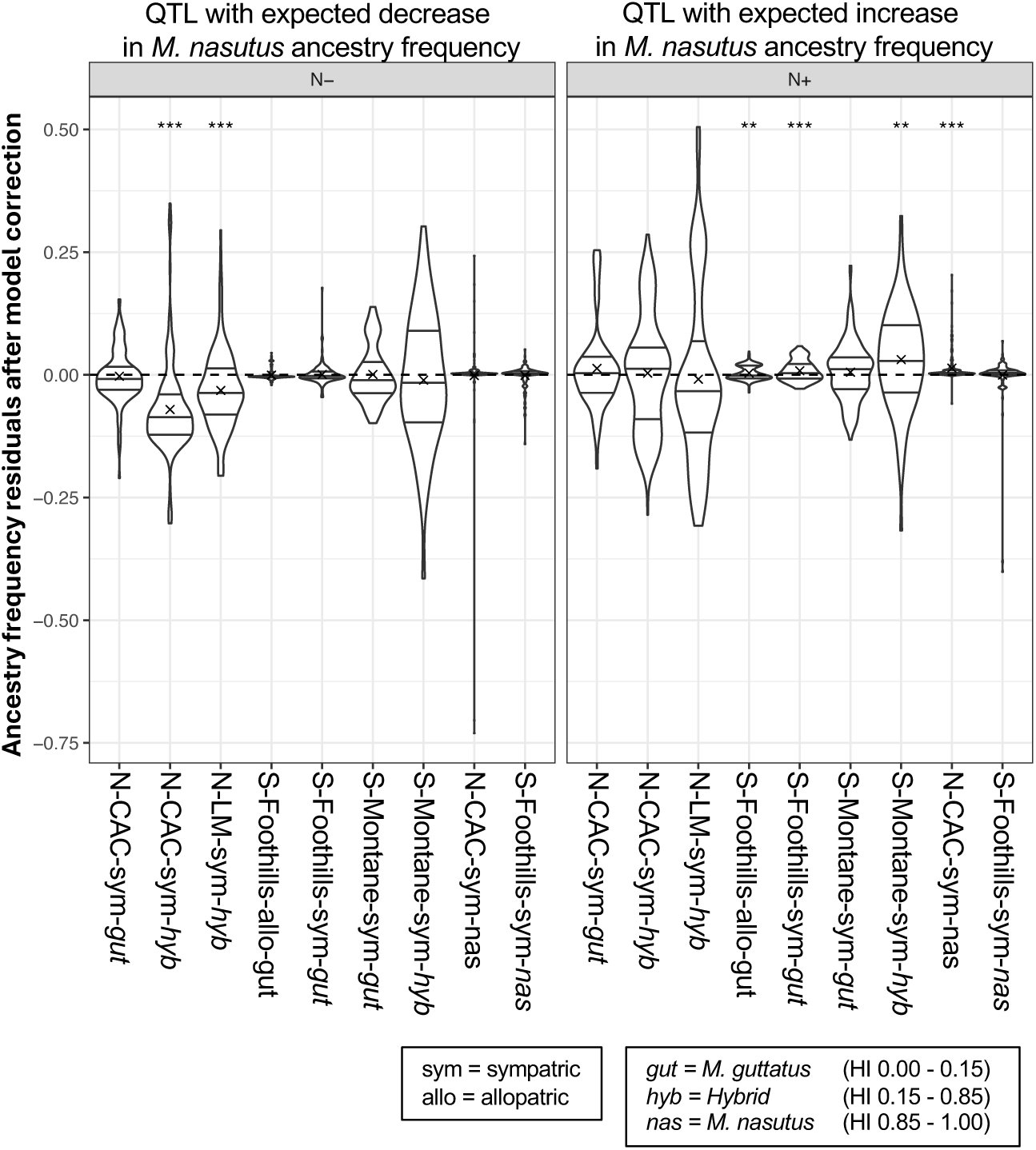
Deviations from expected ancestry frequency for genomic windows in quantitative trait locus regions. Each violin shows the distribution of ancestry frequency residuals, the difference between the observed ancestry frequency and its predicted value from a linear model of genomic features and missingness, for windows contained in a set of quantitative trait loci for intrinsic reproductive isolation as previously identified using crosses from the Northern Catherine Creek population. The ‘N-’ set includes 7 loci with a predicted decrease in *M. nasutus* frequency (*i.e.,* genotype combinations involving N alleles are detrimental), while the ‘N+’ set includes 13 loci with a predicted increase in *M. nasutus* frequency (*i.e.,* genotype combinations involving G alleles are detrimental). For each locus set in each sampling group, we used one-tailed t-tests to mark distributions significantly different from an average deviation of 0 (***p<0.001 and **p<0.01 after Bonferroni correction for 18 tests) in the expected direction.

We predicted that windows within N- QTL would be under-represented among top-5% outliers for ancestry, while N+ QTL windows would be over-represented among top-5% outliers. These patterns were inconsistent across groups, but some groups showed patterns in the expected direction, primarily in the Northern region (Table 2). We found that N- QTL were strongly excluded from ancestry outliers in our three Northern *guttatus* and *hybrid* groups, as well as one Southern group (S-Montane-sym-*guttatus*), but not for other Southern groups or for *nasutus*. Correcting ancestry outliers for genomic structural features and missingness slightly reduced this signal, leading to significant under-representation in only two groups. For N+ QTL, we found no over-representation compared to the uncorrected outliers. After correction for genomic structure and missingness, two groups had a significant overlap with ancestry outliers: CAC *guttatus* and Southern sympatric *guttatus*.

**Table 2.**
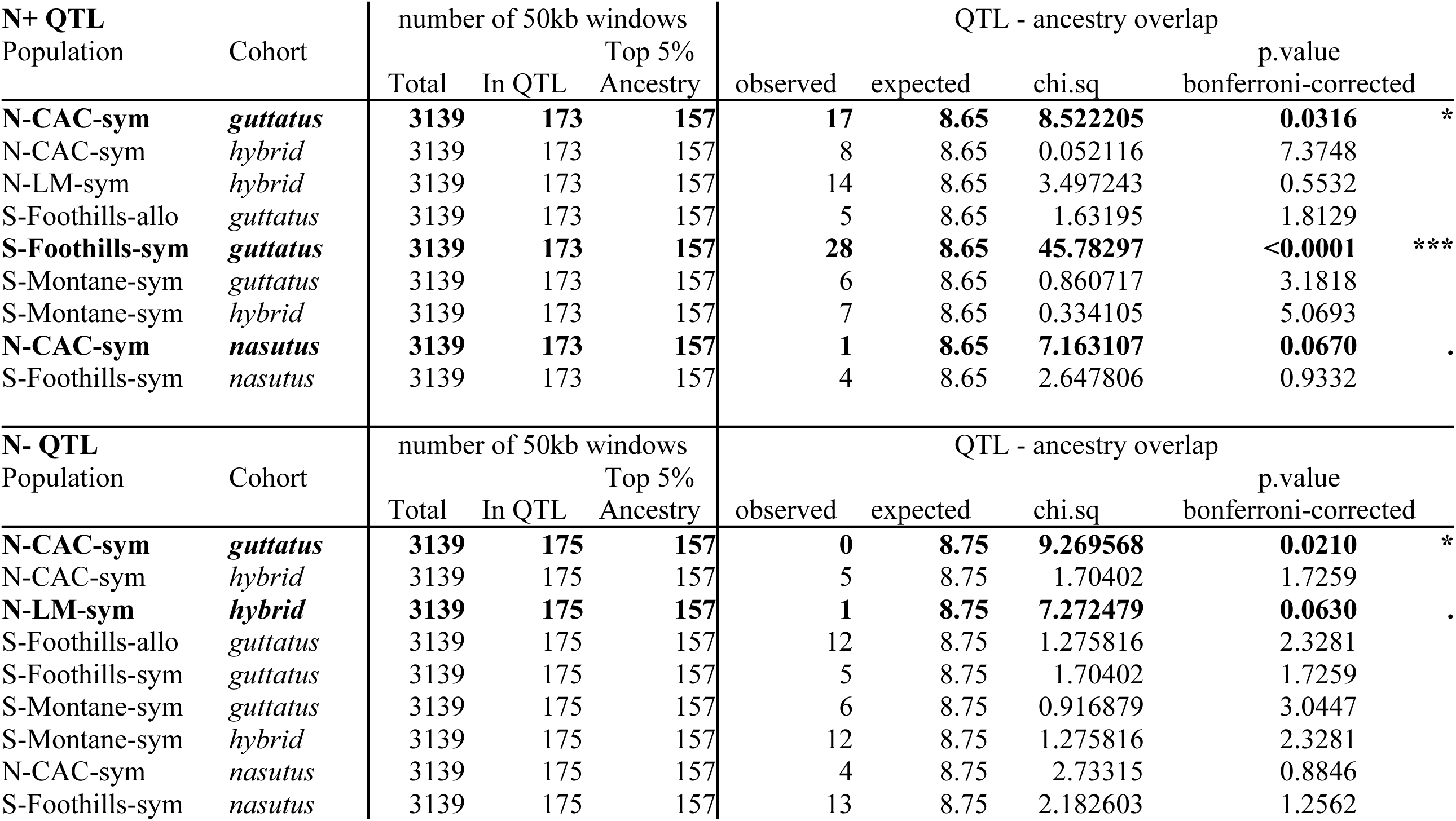
QTL overlap with model-corrected ancestry outliers. Expected counts are based on random selection of windows from the total. N+ QTL = predicted increase in M. nasutus ancestry, N- QTL = predicted decrease in M. nasutus ancestry.

We also examined the genomic regions surrounding four loci known to cause reproductive isolation in some interspecific crosses between *M. guttatus* and *M. nasutus*: two are involved in photoperiod divergence, on Chr07 and Chr08, and two that interact to cause hybrid seedling lethality, on Chr13 and Chr14 (Figure 6). We identified regions of reduced *M. nasutus* ancestry frequency surrounding all four of these loci in at least some groups. However, these loci were far from the most extreme ancestry deviations on their respective chromosomes, suggesting that known loci are, at most, just one of many factors influencing ancestry frequencies. In addition, patterns were often inconsistent across groups, with the same locus at low relative frequency in one group and high relative frequency in another – perhaps not surprising given that *M. guttatus* is known to be polymorphic for functional alleles at all four loci (Fishman et al., 2014; Zuellig & Sweigart, 2018a).

**Figure 6.**
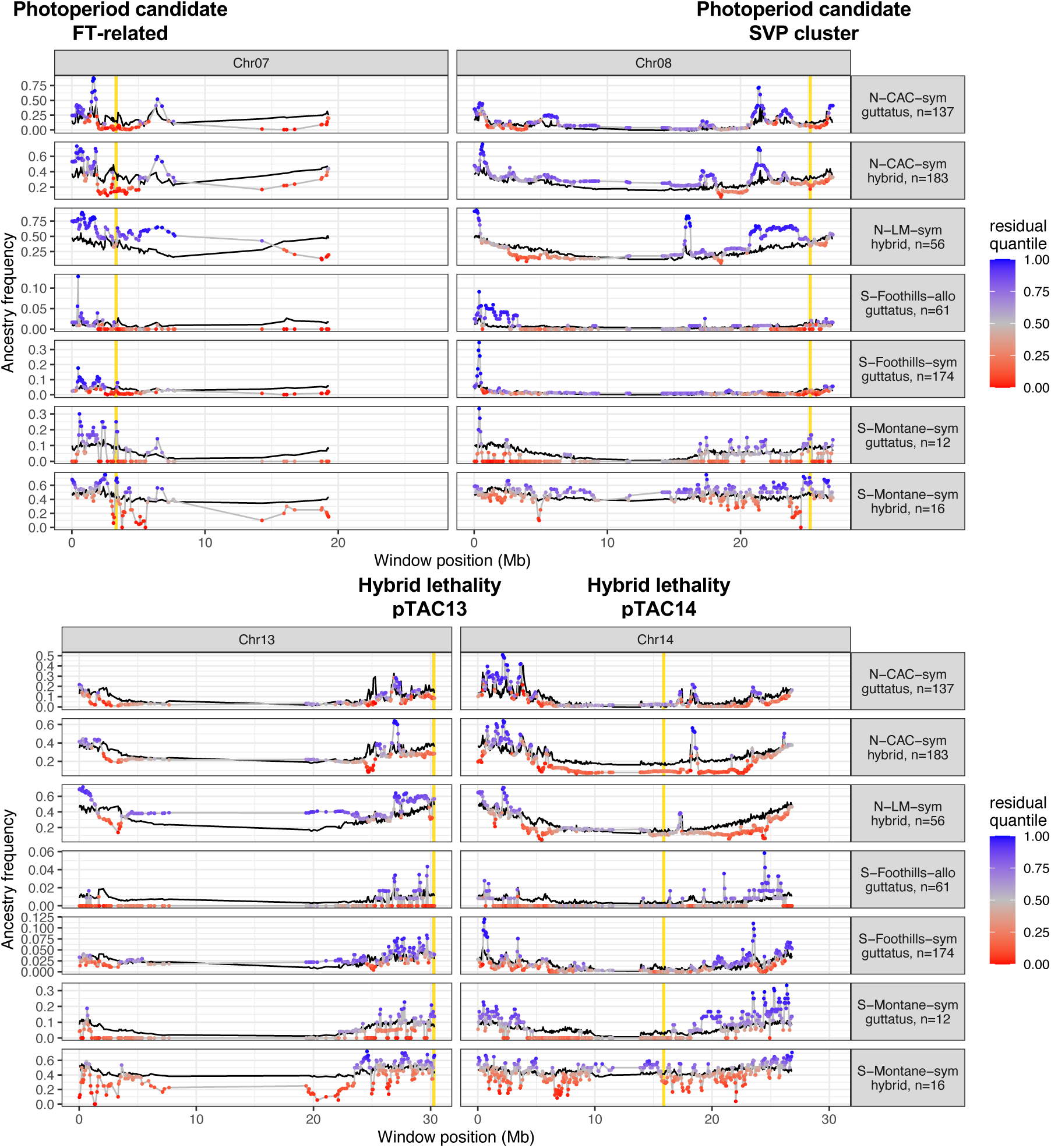
Ancestry frequencies near four known reproductive isolation candidate loci show mixed signals of deviation from genome-wide expectations. Ancestry frequencies (colored points and grey line) vs. predictions from genomic structure and missingness modeling (black line) for sampling groups, focusing on four chromosomes containing known candidate loci involved in *M. guttatus-M. nasutus* reproductive isolation. Candidate locus positions are highlighted in yellow. Colors indicate the empirical quantile within the genome-wide distribution of residual ancestry frequency (i.e., the difference between observed ancestry frequencies and model predictions). *M. nasutus* groups are not shown because the minimal variation in frequency across most of the genome made model predictions and quantile analysis not meaningful in those groups; at all four loci, *M. nasutus* frequencies in *M. nasutus* groups were >0.98.

Our most striking finding was at the Chr07 photoperiod locus, where residual ancestry frequencies were below the bottom 5^th^ percentile for the Northern-CAC *guttatus* and *hybrid* groups (where the photoperiod association was initially characterized (Fishman et al., 2014)) as well as below the bottom 10^th^ percentile for the Southern Foothills sympatric and allopatric *guttatus* groups (Figure 6, Table S5); CAC groups in particular appear to have a strong visual ‘dip’ in ancestry surrounding this locus. However, this same locus was surprisingly above the 95^th^ percentile for residual ancestry in the Southern Montane *guttatus* and *hybrid* groups, though there are large fluctuations among nearby windows in this group. The second photoperiod locus on Chr08 showed less extreme but qualitatively similar patterns, with well-defined ‘dips’ in ancestry in the CAC groups (Figure 6, Table S5). While neither locus was in a particularly low residual quantile in the Northern-LM *hybrid* group, visual ‘dips’ in ancestry relative to surrounding regions were apparent here as well.

At the Chr13 seedling lethality locus, *M. guttatus* segregates for the detrimental allele (Zuellig & Sweigart, 2018b), which if present, should decrease in frequency under hybridization, favoring the protective *M. nasutus* allele instead. However, we see an increase in *M. nasutus* ancestry only in the Southern Montane groups, and instead find a decrease in some Southern Foothills and Northern groups (Figure 6, Table S5); note that the incompatible Chr13 allele has been documented at intermediate frequencies in both the Southern Montane and Southern Foothills area, but its presence in the Northern populations is not known (Zuellig & Sweigart, 2018a). Meanwhile, at the companion locus on Chr14, the detrimental allele is derived from *M. nasutus* (Zuellig & Sweigart, 2018b), so if it is segregating, we would expect a reduction in *M. nasutus* frequency. This expected pattern was again only seen in the Southern Montane *hybrid* group, showing no obvious pattern in other groups (Figure 6, Table S5). However, actual ancestry frequencies near the Chr14 locus were generally low in other Southern and Northern groups; this locus lies in a low-recombination area where predicted ancestry is also low, potentially explaining why it does not appear as an outlier for residual ancestry.

## DISCUSSION

This study provides the first direct comparison of how introgression patterns in the genome vary across populations within an iconic speciation model system, *Mimulus guttatus* vs. *Mimulus nasutus*. We find that A) introgression amounts, symmetries, and genomic distributions vary widely across geographic and genomic space, indicating that a variety of ecological, genomic, and historical processes interact to shaping the outcomes of hybridization; B) shared patterns of ancestry across populations are driven by a combination of genomic landscape features and locus-specific patterns; C) interpopulation gene flow moves introgressing alleles across the landscape of a mosaic hybrid zone; and D) candidate loci from mapping studies are only a partial and inconsistent predictor of natural introgression patterns.

### Wide variation in hybridization outcomes across geographic space

Our data show that hybridization plays an important role in shaping variation throughout the broadly overlapping ranges of *M. guttatus* and *M. nasutus*, with a mosaic of hybrid swarm-like populations, populations with occasional or residual introgression, and even evidence for introgressed alleles in allopatry. This finding is particularly interesting in light of evidence for strong premating (Fishman et al., 2014; Mantel & Sweigart, 2019; N. H. Martin & Willis, 2007) and even some postmating reproductive barriers (Fishman et al., 2008; Fishman & Willis, 2006; Mantel & Sweigart, 2024; N. H. Martin & Willis, 2010; Sweigart et al., 2006; Zuellig & Sweigart, 2018b) between the two species. When taken together, these findings are consistent with an emerging general model of speciation with gene flow during secondary contact, whereby hybridization is difficult to prevent entirely in sympatry (Galloway et al., 2020; Irwin, 2020; Sambatti et al., 2012), but can also persist without completely breaking down species barriers (Feder et al., 2012; Jónsson et al., 2014; Kopp et al., 2018; S. H. Martin et al., 2013; Sianta et al., 2026). Variation across space in the extent of hybridization is a key feature of this model and appears to be common to many hybrid systems (Aboim et al., 2010; Borge et al., 2005; Perrier et al., 2013; Sianta et al., 2024; Simon et al., 2021), including across the *Mimulus guttatus* species complex (Farnitano et al., 2025; Frayer et al., 2026; Tataru et al., 2023). Mosaic hybrid zones, where two species are shaped by divergent selective pressures that vary across a heterogeneous landscape, appear to be particularly prone to this mode of speciation (Larson et al., 2013; Rand & Harrison, 1989; Riquet et al., 2019; Simon et al., 2021).

Surprisingly, we found a large relative number of early-generation hybrids in our Southern Montane group compared to other regions. At these higher elevations, the timing of life-cycle cues – such as snowmelt, temperature, photoperiod, pollinator activity, and water availability – may be very different from other populations (Blackman, 2017; Bucher & Römermann, 2020; Cornelius et al., 2013; Du et al., 2020). We know these cues vary from year to year in Northern populations, impacting hybridization rates (Farnitano et al., 2025); conditions may have been unusually favorable for hybridization in the Montane region in the years leading up to our collections. Variation in the strength of both pre- and post-mating reproductive isolation across time is likely a key contributor to the persistence of hybridizing lineages (Dittmar & Schemske, 2023; Sianta et al., 2024; Tataru et al., 2023).

### Genomic position predicts shared hybridization patterns, but linked selection may not be the reason

Across the genome, we find extreme peaks and troughs of ancestry frequency, both in locations with large hybrid populations and in populations with more limited hybridization, indicating that ancestry patterns are subject to a variety of contrasting forces along the genome. Much of this variation can be attributed to a suite of intercorrelated genomic features: recombination rate, gene density, and chromosomal position. An association between introgression and recombination rate is consistent with earlier findings in *Mimulus* hybrids (Brandvain et al., 2014; Kenney & Sweigart, 2016) as well as many other hybrid systems (Burri et al., 2015; S. H. Martin et al., 2019; Nouhaud et al., 2022; Schumer et al., 2018), where they have been taken as evidence for pervasive selection against minor-parent ancestry. However, when we use a more direct proxy for the effects of linked selection (gene density scaled by recombination rate), we see no association with minor-parent ancestry, suggesting that linked selection is not directly responsible for these patterns. This is consistent with recent findings of weak linked selection operating within *Mimulus guttatus* (Lovell et al., 2025) and other plant species (Fernandes et al., 2024; Slotte, 2014), but to our knowledge this lack of a pattern has not been remarked upon in hybridizing populations before now.

If linked selection against hybrid ancestry is not driving ancestry patterns, then why is minor-parent ancestry consistently higher towards the ends of chromosomes? One possibility is that a form of linked selection is still effective in the earliest stages of hybridization, when chromosomes are broken up by only one or a few crossovers; this could cause these very-low-recombination regions in the center of chromosomes to be purged, while large chunks of chromosome arms are retained. As more recombination events occur in later-generation hybrids, the efficacy of linked selection may rapidly diminish so that no finer-scale correlation is apparent. Another possibility is that a mixture of positive and negative selection on introgressing alleles cancel each other out in relatively gene-dense regions, but gene-poor regions are dominated by negative selection. Further theoretical and empirical work is needed to investigate the reasons for a lack of linked selection associations in plant species, and what other forces are shaping these genomic patterns in plant hybrids.

### Introgressed haplotypes are transported from sympatry to allopatry

Geographically closer groups tend to have more similar ancestry frequencies, even after accounting for genomic features and missingness, consistent with a role of gene flow spreading alleles from their source of introgression across the mosaic landscape. In particular, the small amounts of introgression in allopatric Southern Foothills populations are best explained by introgression patterns of nearby sympatric populations, suggesting they may have received these alleles through interpopulation gene flow. Another possible explanation of ancestry correlations in nearby populations is greater similarity in their genomic background leading to parallel selection patterns. While these genomic background effects may be important, they alone would not be expected to produce any introgression in allopatry, so our findings still support gene flow as an important source of correlated introgression. Other studies have shown that introgressed alleles can filter out into the wider allopatric population (Martinsen et al., 2001; Sankararaman et al., 2014; Simon et al., 2021), and even become fixed (Nelson et al., 2021; Pardo-Diaz et al., 2012), though the extent and speed at which this happens will vary depending on demographic conditions and the strength of selection (Piálek & Barton, 1997; Sachdeva & Barton, 2018). Percolation through multiple steps of migration is likely to selectively filter only the most adaptive (or at least the least deleterious) introgressing loci, providing an important source of genetic variation for the species as a whole (Baskett & Gomulkiewicz, 2011; Oziolor et al., 2019); this concept has been used effectively to reveal adaptive introgression candidates (Cruzan et al., 2021; Gaczorek et al., 2024; Gompert & Buerkle, 2011; Sung et al., 2018). Our findings support the conclusion that the evolutionary impacts of hybridization are not restricted to narrowly defined hybrid zones, but can have substantial impacts on species-wide trajectories (Mallet, 2005; Taylor & Larson, 2019).

### A detectable but limited role for locus-specific parallel selection in shaping hybridization patterns

After other sources were accounted for, we found a reduced, but detectable, pattern of residual shared variance in ancestry between sympatric groups in the Northern and Southern regions. Unlike the allele sharing between nearby populations, this pattern is unlikely to be caused by gene flow given the large geographic distance (∼1000km) and substantial genetic differentiation between Northern and Southern *M. guttatus*. Instead, we attribute this pattern to parallel, but independent, selection on the same genomic loci across both regions. Either positive (favoring introgression) or negative (resisting introgression) selection pressures could be contributing to these patterns; in fact, the wide range of ancestry frequencies across the genome suggests that both are at play. Large swings from high to low ancestry across the genome are particularly apparent in the Northern *hybrid* cohorts, consistent with a ‘hybrid bridge’ hypothesis: an ongoing hybrid population provides an opportunity for adaptive alleles to become unlinked from introgression-resistant regions and more readily cross species boundaries (Martinsen et al., 2001; Simon et al., 2021). We investigated the sharing of top introgression outliers and found significant overlap across groups even after genomic features correction; these overlapping outliers are promising candidates for future exploration as targets of adaptive introgression.

### Known reproductive isolation loci can reduce introgression, but patterns are often population-specific

Our tests of ancestry patterns surrounding reproductive isolation loci revealed some evidence that these loci were resistant to introgression, but patterns were highly population-specific. The 20 RIL-derived QTL (Mantel & Sweigart, 2024) were originally detected in crosses derived from CAC individuals; one explanation is that the particular alleles present in those crosses are geographically localized and not found in the South. Geographically localized reproductive isolation loci are a common pattern between these species (Mantel & Sweigart, 2024; N. H. Martin & Willis, 2010; Sweigart et al., 2007; Zuellig & Sweigart, 2018a), fitting with theory suggesting that polymorphic reproductive isolation is a common and influential feature in early stages of speciation (R. B. Corbett-Detig et al., 2013; Cutter, 2012; Servedio & Hermisson, 2020). Detecting a deficit of introgression is also statistically difficult in populations where the majority of the genome already has low or no introgression. Also, the wide confidence intervals from these QTL are probably much larger than the scale of linkage in natural hybridizing populations (Brandvain et al., 2014; Puzey et al., 2017), so reduced introgression at the causal locus may be diluted by nearby windows. Finally, these QTL likely represent only a small sampling of loci under selection in wild populations, especially if the genetic basis of reproductive isolation is highly polygenic. Therefore, any signals at these loci may not be outliers compared to the average genomic pattern. Together, these limitations highlight the challenges in making direct comparisons between trait-associated, lab-based studies of reproductive isolation and the complex patterns that appear in natural hybridizing populations.

Two large-effect photoperiod candidate loci (Fishman et al., 2014) did appear to have an effect on introgression patterns, but again, the patterns were population-specific. Interestingly, the Southern Montane populations showed a pattern opposite to other *guttatus* groups, with high apparent introgression at these loci. This fits with our earlier observation that Montane populations may have a different relationship to life cycle timing than lower-elevation groups; photoperiod alleles may in fact be introgressing adaptively in this region rather than serving as a reproductive barrier. However, photoperiod as a trait is highly variable across the large and ecologically variable range of *M. guttatus* (Friedman & Willis, 2013; Monnahan & Kelly, 2017), and indeed the underlying loci are likely polymorphic (Zuellig et al., 2014), so in different populations we might be detecting the effect of different alleles entirely. These patterns again highlight how environmental variation can drive differences among populations in the extent and genomic targets of introgression.

Two known interacting hybrid lethality loci on Chr13 and Chr14 (Zuellig & Sweigart, 2018b) were expected to be strong candidates for introgression outliers, but their patterns matched our expectations only in the Southern Montane region, and in fact Chr13 showed patterns opposite to our expectations in some other groups. These loci are known to be polymorphic rather than fixed across both species (Zuellig & Sweigart, 2018a), so some of this inconsistency may be due to other non-lethal alleles taking precedence in certain populations. The interacting haplotypes are also both recessive, which might limit the efficacy of selection especially in groups where new hybridization is rare. Finally, there may not be strong positive selection on introgression for the ‘protective’ alleles even if there is negative selection against introgression of the incompatible alleles, since only one protective allele in the pair is necessary for viability. While extensive theoretical work has been devoted to predicting the effects of such classical incompatibilities (Bank et al., 2012; Lemmon & Kirkpatrick, 2006; Lindtke & Buerkle, 2015; Ravinet et al., 2017), our data show that the realities of population variability and polymorphism quickly complicate those effects in real populations (Cutter, 2012).

Overall, our analysis suggests that outlier loci detected via ancestry patterns in natural populations are often complementary to, rather than redundant with, reproductive isolation loci detected by trait mapping approaches. Most of the outlier regions we find, even those shared among populations, are not tied to known pre- or postmating loci. Presumably at least some of these have important selective effects in admixed populations, though others may be the result of drift or be affected by directional selection unrelated to species differences. We stress that neither trait mapping nor admixture scanning is sufficient to fully understand the genetic basis of species barriers; instead, we can learn a great deal by unifying the findings from multiple independent approaches. Some studies have used introgression patterns to detect or confirm candidate loci from mapping studies (Larson et al., 2013; Lindtke & Buerkle, 2015; Rifkin et al., 2019). But other recent work (Feller et al., 2024; Frayer & Payseur, 2024) has shown a lack of correspondence between natural introgression patterns and laboratory-based trait mapping studies. Among other explanations, they argue that the differing time scales of ongoing contact in hybrid zones vs. direct crosses are partly responsible. Our study shows this discrepancy may be common, and points to some additional potential reasons: polymorphism across geographic space, population-specific selection patterns, highly polygenic architectures of reproductive isolation, and differing resolutions of QTL vs. admixture outliers.

### Technical factors may affect the identification of shared hybridization patterns

Certain regions of the genome may be prone to misidentification during the ancestry calling process, for both biological and technical reasons. We found that windows with higher proportions of missing data also tend to have higher minor-parent ancestry, and that this can explain a substantial proportion of ancestry variation. We consider three possible reasons for such a correlation, though there may be others. First, certain regions might have a read-mapping bias that favors the minor parent; this could simultaneously cause higher amounts of missing data and an erroneously high minor-parent ancestry. In this case, the association is purely technical and should not be attributed to biological causes. Read-mapping bias is more likely to favor *M. nasutus* ancestry because of the much higher structural and nucleotide diversity present within *M. guttatus* (Brandvain et al., 2014; Lovell et al., 2025). Second, incomplete lineage sorting, which is common due to the nested nature of *M. guttatus* - *M. nasutus* divergence (Brandvain et al., 2014; Lovell et al., 2025), might cause certain variants, which are in fact segregating within *M. guttatus* prior to introgression, to be used as evidence for *M. nasutus* ancestry; these variants would also provide conflicting signals to other nearby variants, leading to lower genotyping confidence in the region. Third, certain genomic features, such as low selective constraint or high levels of structural variation, might lead to both higher actual introgression (due to reduced selection against minor-parent ancestry) and higher levels of missingness (due to poor read mapping or a lack of informative markers). In this final case, our association with missingness would represent a true biological signal rather than a technical limitation, and would be an intriguing pattern worthy of future study. There is little data currently available on the expected impacts of highly-variable regions on introgression outcomes, but a few studies point to an important role for structural variation in introgressed regions (Hsieh et al., 2025; Upadhyay et al., 2021; Zhang et al., 2023). Distinguishing between these hypotheses will require further high-coverage sequencing and new ancestry inference tools to reconstruct the ancestry of discrete haplotypes with high confidence (Huang et al., 2025). Moving forward, we recommend that researchers in other systems pay close attention to the effects of genotyping or ancestry-calling uncertainty on their interpretation of genome-wide patterns.

### Conclusion

Overall, our system provides a strong example of an ongoing and dynamic hybridization mosaic without the collapse of species boundaries. Furthermore, our results argue for a combination of both universal, repeatable patterns and idiosyncratic population- and locus-specific effects shaping the genomic landscape of hybridization. However, the subtler details of these patterns argue for caution and nuance in how we conventionally interpret genomic signals of hybridization. We find a lack of evidence for linked selection in shaping the genomic landscape, and only minor or inconsistent effects of key ‘speciation genes’ from mapping studies. The field of hybridization genomics needs to work to better understand the root causes of variation across the genome in hybrid ancestry, through a combination of careful comparative work across systems, robust theory and modeling, and a better understanding of the multidimensional landscape of selection operating on hybrids.

## ACKNOWLEDGEMENTS

We thank members of the Sweigart Lab, including Eleanore Ritter, Natalie Gonzalez, Logan Scott, and Lauren Womack, for comments on drafts of this manuscript. Molly Schumer, Kelly Dyer, Casey Bergman, Jill Anderson, Keith Karoly, John Willis, and John Kelly provided guidance and advice during the development of this project. Sara Hill, Autumn Knight, Liza Lesley, Sherwin Shirazi, and Parker Helms assisted with sample processing for sequencing. We also thank Bob Schmitz for providing the tagmentation enzyme for library preparation.

## DATA AVAILABILITY STATEMENT

All Illumina data generated for this project will be archived at the Sequence Read Archive (SRA) upon manuscript acceptance. Ancestry calls, sample metadata, and other processed data will be made available on the DRYAD digital repository. All analysis scripts are available on GitHub (github.com/mfarnitano/Mimulus_hybrid_ancestry).

## BENEFIT-SHARING STATEMENT

All samples collected for this project were done so in accordance with local laws and with the cooperation of local agencies.

## AUTHOR CONTRIBUTIONS

MCF designed the project, collected the data, conducted the analysis, and wrote the paper. VAS helped to design the project and collect the data. ALS helped to design the project, revised and edited the paper, and secured funding.

## SUPPLEMENTAL FILES

### Supplementary Text

1. Genotyping methods details.
2. Excluding windows with poor fit of ancestry to local PCA.
3. Splitting of allopatric samples into high- and low-introgression subgroups.
4. Supplemental Text References.

### Supplementary Tables

Table S1. Sample sizes before and after filtering across all groups.

Table S2. *M. guttatus* and *M. nasutus* individuals used in an ancestry reference panel.

Table S3. Relationship between ancestry and genomic features before and after excluding pericentromeric regions.

Table S4. Breakdown of *M. guttatus* (HI<0.15) individuals within Southern populations.

Table S5. Raw and residual ancestry frequency quantiles at four fine-mapped reproductive isolation loci.

Table S6. Grouping of ancestry window outliers into contiguous stretches.

### Supplementary Figures

Figure S1. Distributions of R^2^_PC1_ and |Z_het_| scores for northern and southern regions.

Figure S2. Histograms of hybrid index recalculated with reduced window dataset.

Figure S3. Ancestry frequency distributions recalculated with reduced window dataset.

Figure S4. Proportion of variance in ancestry frequency explained by genomic features, missingness, and other sample groups, recalculated with reduced window dataset.

Figure S5. Overlap among ancestry frequency outlier windows across groups, after correction for genomic features and missingness, recalculated with reduced window dataset.

Figure S6. Residual ancestry frequency distributions in reproductive isolation QTL regions, recalculated with reduced window dataset.

Figure S7. Distance to the nearest sympatric location is negatively correlated with average hybrid index of allopatric Foothills locations.

Figure S8. Proportion of variance in ancestry frequency explained by genomic features, missingness, and other sample groups, with the Southern Foothills allopatric split into allopatric-low and allopatric-high subgroups.

Figure S9. Overlap among ancestry frequency outlier windows across groups, after correction for genomic features and missingness, with the Southern Foothills allopatric group split into allopatric-low and allopatric-high subgroups.

Figures S10-S13. Ancestry frequencies compared to model predictions for all 14 chromosomes.

