## Supplementary Text for "On the causes of correlated genomic ancestry across contrasting hybridization histories in a monkeyflower species pair"

This supplementary text accompanies the manuscript "On the causes of correlated genomic ancestry across contrasting hybridization histories in a monkeyflower species pair" by Farnitano, Matthew C., V. Alex Sotola, and Andrea L. Sweigart.

#### I. Genotyping methods details.

To genotype our reference panel of 33 high-coverage inbred lines, we obtained paired-end fastq files, randomly downsampled three very-high-coverage samples to 60 million read pairs, trimmed low-quality ends with *Trimmomatic v0.39* (Bolger et al., 2014), aligned reads to the *M. guttatus* IM767 v2 reference assembly (Lovell et al., 2025) using *bwa v0.7.17*(Li & Durbin, 2009), removed duplicates with *picard v2.27.5* (Broad Institute, 2019), filtered for properly-paired reads with MAPQ>=29 using *samtools v1.16.1* (Danecek et al., 2021), then called SNPs with *GATK v 4.4.0.0 HaplotypeCaller* (Van der Auwera & O'Connor, 2020). To obtain a list of high-quality SNPs, we used *GATK* to filter for biallelic sites that met the following criteria: QD>=2, QUAL>=40, SOR<=3, FS<=60, MQ>=40, MQRankSum >= -12.5, ReadPosRankSum >= -12.5, and ReadPosRankSum <= 12.5, as well as restricting to only sites within annotated genes and outside of repeat-masked regions of the IM767 reference genome. We then set individual sample genotypes to missing if they had a GQ<15 or an individual DP<6 or DP>100.

To assess population structure and variation across northern and southern samples, we used *angsd v0.940* (Korneliussen et al., 2014) with the GATK likelihood method to estimate genotype likelihoods at each *angsd* marker for each individual. Low-coverage samples may be sensitive to false homozygosity if overlapping forward and reverse reads are counted twice during genotype likelihood estimation, so we first merged overlapping forward and reverse reads using *Flash v 2.2.00* (Magoč & Salzberg, 2011). Then, we trimmed, mapped, and filtered reads using the same approach and parameters as the reference panel above, and used the resulting alignments as input for *angsd* genotype likelihood estimation.

#### II. Excluding windows with poor fit of ancestry to local PCA.

***Methods.*** One potential source of error in ancestry assignments is genetic structure that does not correspond with interspecific ancestry, because of either incomplete lineage sorting, geographical structure, or other historical patterns. The substantial genetic divergence between northern and southern *M. guttatus* (see PCA in Main text Figure 1) in particular complicates ancestry assignment. We therefore ran local PCAs using genotype likelihoods for each individual 50kb genomic window, in order to test the correspondence between local population structure and inferred species ancestry. We ran separate PCAs for the northern region and for the southern region (excluding LM samples and high-elevation southern populations for simplicity), under the assumption that the primary axis of variation within each region should be *M. guttatus – M. nasutus* ancestry. Then, we calculated two measures of the degree of fit between these local PCAs and our HMM-based ancestry assignments. First, we calculated the coefficient of determination ($R^2_{PC1}$) between the first local PC axis and the windowed ancestry assignment. Second, we calculated a Z-score ($|Z_{het}|$) representing the degree to which individuals with a heterozygote ancestry assignment deviate from the expected midpoint between alternative ancestry homozygotes along the first local PC axis (Equation S1). Distributions of these two measures are shown in Figure S7. We created a 'high-confidence reduced window set' including only genomic windows with $R^2_{PC1}$ >=0.9 and $|Z_{het}|$ <= 2 for both the northern region and southern region sample groups. We re-ran analyses of ancestry correlations and outlier window analyses using only this reduced set.

***Equation S1.*** Deviations of $|Z_{het}|$ from 0 indicate that heterozygous samples on average fall off of the midpoint between homozygous *guttatus* and homozygous *nasutus* samples on PC1.

$u_G$ = mean(PC1) for windows with ancestry genotype=*guttatus homozygous*
$u_H$ = mean(PC1) for windows with ancestry genotype=*heterozygous*
$u_N$ = mean(PC1) for windows with ancestry genotype=*nasutus homozygous*
$u_{exp} = (u_G - u_N)/2$
$u_{scaled} = (PC1 - u_{exp})/(u_N - u_{exp})$ for windows with ancestry genotype=*heterozygous*
$|Z_{het}| = |\ mean(u_{scaled})/sqrt(var(u_{scaled}))\ |$

***Results.*** We excluded a total of 1071 windows (34%) with low $R^2_{PC1}$ or high $|Z_{het}|$ values, leaving a remaining 'reduced' dataset of 2069 windows (Figure S1). Notably, this excluded a handful of the most extreme ancestry outlier windows for both *M. nasutus* groups and for the Southern sympatric *M. guttatus* group, but overall hybrid index distributions and ancestry frequency distributions were not strongly affected (Figure S2-S3). Overall, ancestry correlations across groups were lower for this reduced dataset, both before and after genomic structure and missingness corrections, but still significant for many comparisons and consistent with the general patterns in the full dataset (Figure S4). In particular, correlations between other groups and the Southern Foothills allopatric group were much lower, suggesting some of these correlations may be driven by the lower-confidence windows that were removed from this reduced dataset. However, some correlations with allopatry do remain, including a significant correlation between Southern Foothills allopatric and sympatric *guttatus* groups (Figure S4).

There was also substantially less overlap among ancestry outliers across groups for this reduced dataset, especially comparing Southern to Northern groups (Figure S5). This suggests that many of the shared outlier windows in our allopatric and high-elevation set may be windows with poor correspondence between ancestry calls and local PCA. These outliers are more likely to be biased by technical reasons related to poor ancestry assignment, rather than true introgression patterns, though they could still be true cases of outlier introgression in regions with complex evolutionary histories or low read mapping quality. Crucially, however, the Southern sympatric and allopatric groups retain their significant outlier overlap with the Northern CAC *hybrid* group, indicating that this cross-region signal in sympatry persists in at least some comparisons, despite stringent filtering procedures. Also, the Southern allopatric retains strong overlap with the Southern sympatric group, indicating a set of 17 high-confidence windows with a robust allopatric introgression signal indicative of gene flow from sympatry (Figure S5). Finally, our analyses of QTL overlap with ancestry outliers was not strongly affected by the use of the reduced dataset; a few comparisons that were significant in the full dataset were no longer significant in the reduced dataset, highlighting the weak and population-specific nature of QTL effects on ancestry (Figure S6).

### III. Splitting of allopatric samples into high- and low-introgression subgroups.

***Methods.*** Grouping southern *M. guttatus* (individuals with hybrid index <=0.15) individuals by collection location revealed that some locations carried higher *M. nasutus* admixture than others, even among allopatric populations (Table S1). In particular, two allopatric populations (MOC and RCF) had average hybrid indices greater than 0.01, comparable to sympatric population averages; a third allopatric population (BFR) had one individual with hybrid index >=0.01. We obtained the distance from each allopatric location to the nearest sympatric location in our dataset, using GPS coordinates and the R package 'geosphere.' We then compared this distance to the average hybrid index of the allopatric population using a linear model (Figure S1). In order to examine whether the three populations with higher drive our apparent signals of correlation between allopatric and sympatric *M. guttatus*, we re-analyzed uncorrected and corrected correlations after splitting the 'Southern allopatric' group into two separate subgroups. The first subgroup, 'Southern allopatric-high', included the three populations listed above with at least one individual of hybrid index >=0.01 (MOC, RCF, and BFR; 32 individuals total). The second subgroup, 'Southern allopatric-low', included only allopatric populations with a maximum individual hybrid index <0.01 (three populations: COP, GCH, RHI; 29 individuals total).

We selected the top 5% outlier windows for introgression frequency in these subgroups and compared their frequency of overlap with outliers in other groups, following the same approach detailed in the main manuscript *Methods*. To assess the contiguity of these overlaps, we counted the number of unbroken stretches of outlier windows within the genome for each group and subgroup (Table S2).

***Results.*** Allopatric locations that were geographically closer to nearby sympatric populations tended to have higher mean hybrid indices, consistent with an effect of gene flow from sympatry into allopatry (Figure S7). Overall, the 'allopatric-high' group retained similar correlations with other sympatric populations as the combined allopatric group. In contrast, the 'allopatric-low' group had substantially reduced correlations with northern and high-elevation populations, indicating that the 'allopatric-high' populations were driving most of the allopatric correlation signal. However, even the 'allopatric-low' group retains a substantial correlation with the southern sympatric group (Figure S8). This persistent correlation indicates that even these populations have small amounts of introgression via gene flow from sympatric populations. Alternatively, the remaining correlation could represent a spurious signal due to erroneous ancestry calls at a subset of windows.

Outlier windows in the 'allopatric-low' group are fragmented into more, smaller stretches compared to southern sympatric, allopatric-high, and northern groups, consistent with these windows being older stretches of introgression broken up by more generations of recombination, though this signal could also be indicative of spurious windows (Table S6).

Top outlier windows in the southern 'allopatric-low' subgroup were less likely to overlap with northern group outlier windows than the 'allopatric-high' subgroup (Figure S9). However, even 'allopatric-low' outliers overlapped significantly with Foothills sympatric outliers, consistent with gene flow from sympatry being responsible for the presence of these allopatric-low outliers. The 20 identified windows that are significant outliers for ancestry in both the Foothills-sympatic and Foothills-allopatric-low groups are strong candidates for positive selection acting on introgression in those windows (Figure S9).

### **IV.** Supplemental Text References


Bolger, A. M., Lohse, M., & Usadel, B. (2014). Trimmomatic: A flexible trimmer for Illumina sequence data. *Bioinformatics*, *30*(15), 2114–2120. https://doi.org/10.1093/bioinformatics/btu170

Broad Institute. (2019). *Picard Toolkit, GitHub repository* [Computer software]. https://broadinstitute.github.io/picard/

Danecek, P., Bonfield, J. K., Liddle, J., Marshall, J., Ohan, V., Pollard, M. O., Whitwham, A., Keane, T., McCarthy, S. A., Davies, R. M., & Li, H. (2021). Twelve years of SAMtools and BCFtools. *GigaScience*, *10*(2), giab008. https://doi.org/10.1093/gigascience/giab008

Korneliussen, T. S., Albrechtsen, A., & Nielsen, R. (2014). ANGSD: Analysis of Next Generation Sequencing Data. *BMC Bioinformatics*, *15*(1), 356. https://doi.org/10.1186/s12859-014-0356-4

Li, H., & Durbin, R. (2009). Fast and accurate short read alignment with Burrows-Wheeler transform. *Bioinformatics*, *25*(14), 1754–1760. https://doi.org/10.1093/bioinformatics/btp324

Lovell, J. T., Walstead, R., Lawrence, A., Stark-Dykema, E., Farnitano, M. C., Harder, A., Brůna, T., Barry, K., Goodstein, D., Jenkins, J., Lipzen, A., Boston, L., Webber, J., Chovatia, M., Eichenberger, J., Talag, J., Grimwood, J., Schmutz, J., Kelly, J. K., … Willis, J. H. (2025). Comparative Analyses of Four Reference Genomes Reveal Exceptional Diversity and Weak Linked Selection in the Yellow Monkeyflower (Mimulus guttatus) Complex. *Molecular Ecology Resources*, *25*(8), e70012. https://doi.org/10.1111/1755-0998.70012

Magoč, T., & Salzberg, S. L. (2011). FLASH: Fast length adjustment of short reads to improve genome assemblies. *Bioinformatics*, *27*(21), 2957–2963. https://doi.org/10.1093/bioinformatics/btr507

Van der Auwera, G., & O'Connor, B. (2020). *Genomics in the Cloud: Using Docker, GATK, and WDL in Terra (1st Edition)*. O'Reilly Media.
