## Supplementary Figures for "On the causes of correlated genomic ancestry across contrasting hybridization histories in a monkeyflower species pair"

These supplementary figures accompany the manuscript "On the causes of correlated genomic ancestry across contrasting hybridization histories in a monkeyflower species pair" by Farnitano, Matthew C., V. Alex Sotola, and Andrea L. Sweigart.

**Figure S1. Distributions of $R^2_{PC1}$ and $|Z_{het}|$ scores for northern and southern regions.** See Supplementary Text for description of these values; briefly, $R^2_{PC1}$ indicates the strength of the correlation between local-window genomic PC1 and ancestry genotype assignment, while $|Z_{het}|$ indicates the extent that local-window genomic PC1 values for samples with heterozygous genotypes deviate from the midpoint of average PC1 values for samples with homozygous *nasutus* and homozygous *guttatus* genotypes. Each point represents a 50kb genomic window. Dotted red lines indicate cutoffs for removal of windows to create a high-confidence 'reduced' dataset.

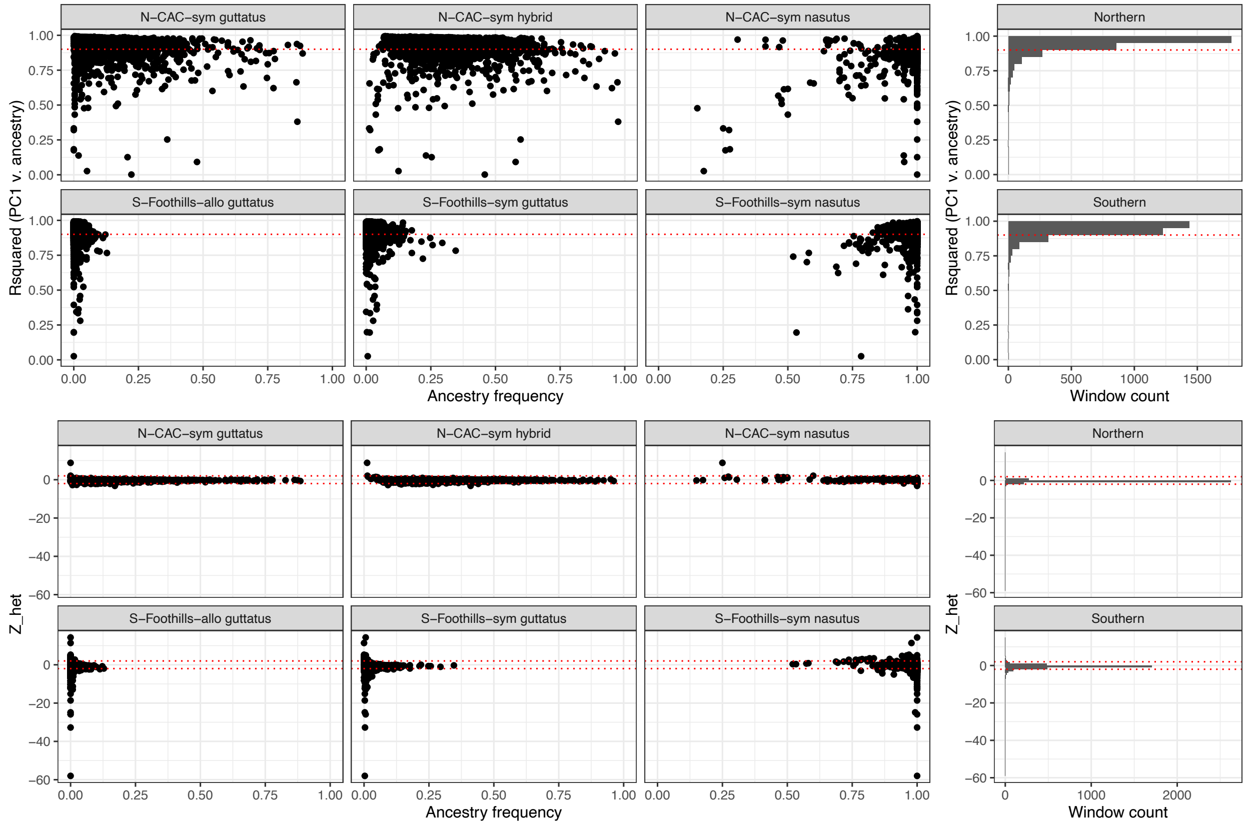

**Figure S2. Histograms of hybrid index recalculated with reduced window dataset.** Compare to Main Text Figure 1E. Removal of lower-confidence windows did not substantially affect the overall distributions of hybrid index for each sample group.

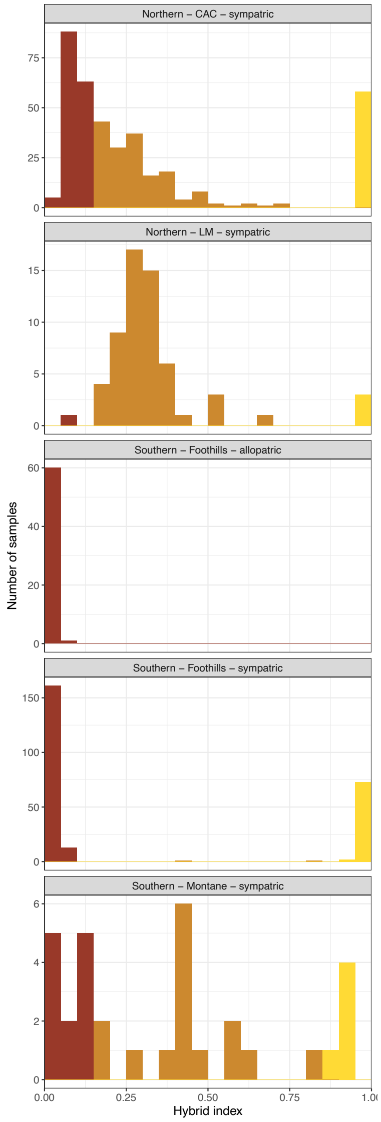

**Figure S3. Ancestry frequency distributions recalculated with reduced window dataset.** Compare to Main Text Figure 2A. Removal of lower-confidence windows had an effect on the most extreme ends of the frequency distribution for some groups, particularly Southern Foothills sympatric *guttatus* and *nasutus*, but otherwise did not strongly affect overall ancestry frequency distributions.

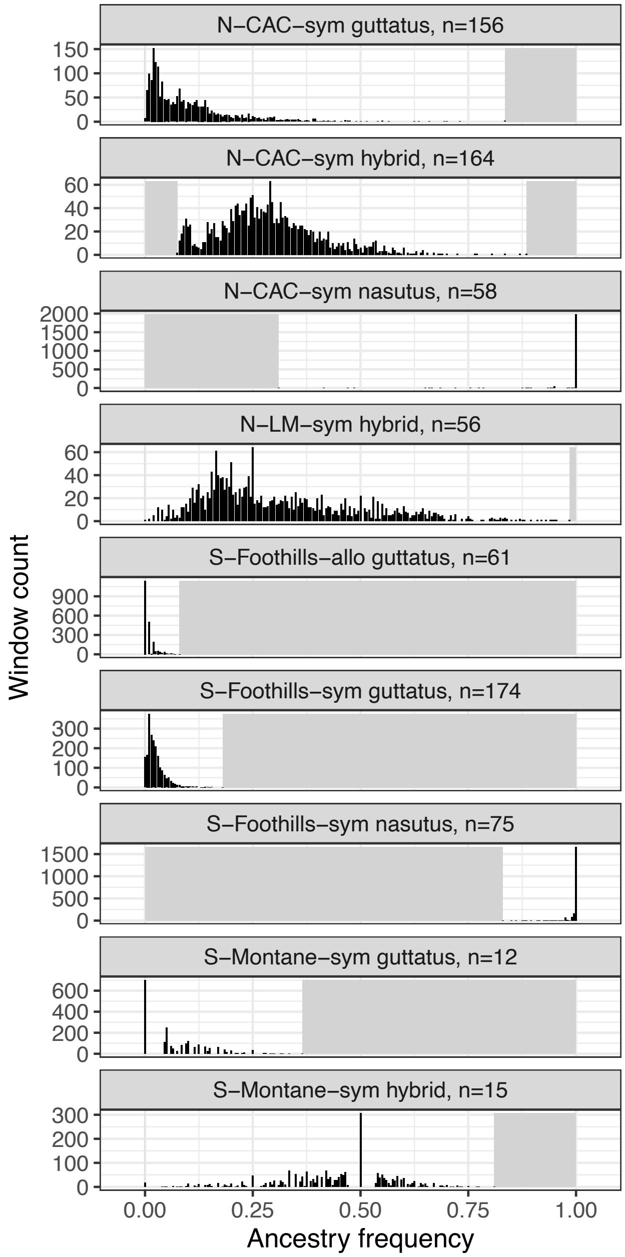

**Figure S4. Proportion of variance in ancestry frequency explained by genomic features, missingness, and other sample groups, recalculated with reduced window dataset.** Compare to Main Text Figure 3A. Note that ancestry correlations overall were lower in this reduced set compared to the full dataset, but the same general patterns were present: stronger correlations among geographically closer groups, including between allopatric and sympatric Southern Foothills groups, and some retained correlations even between Northern and Southern *guttatus* or *hybrid* groups. Higher correlations between sympatric Foothills *guttatus* and other groups, compared to between allopatric Foothills *guttatus* and other groups, suggest that some of the correlations with allopatry in the full dataset may be driven by lower-confidence windows.

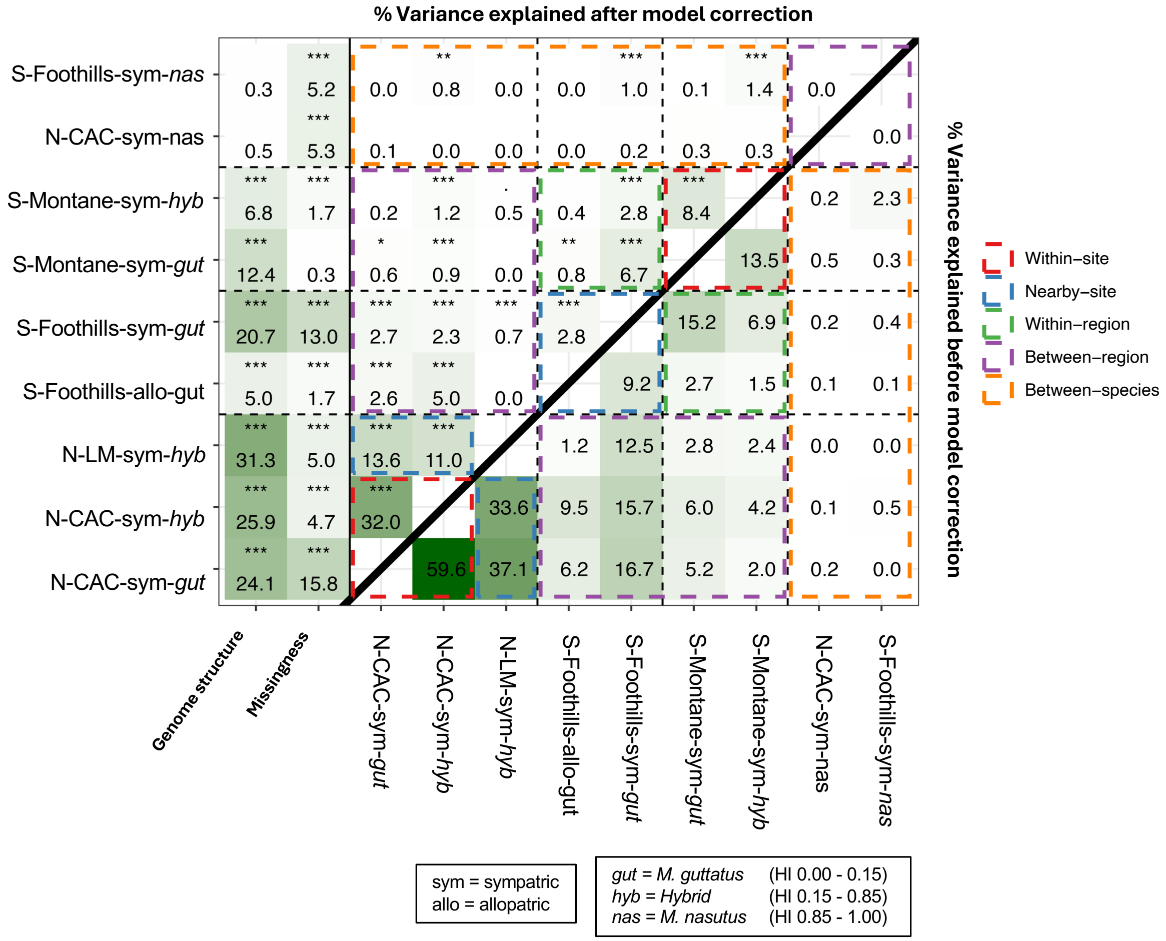

**Figure S5. Overlap among ancestry frequency outlier windows across groups, after correction for genomic features and missingness, recalculated with reduced window dataset.** Compare to Main Text Figure 4A. Overlap among outliers in this reduced dataset was substantially smaller in many pairs compared to the full dataset. However, some significant signals remain which are robust to a stringent choice of genomic windows. There are significant within-region overlaps in the Northern groups as well as between allopatric and sympatric Southern Foothills groups, corroborating our signal of within-region gene flow spreading introgression signals between populations. Note that the uncorrected comparisons involving N-CAC-sym-nasutus are not meaningful because fewer than 5% of windows have ancestry frequencies <1.0 in this group, meaning that all 2069 windows trivially qualify at the 5% cutoff as outliers for introgression.

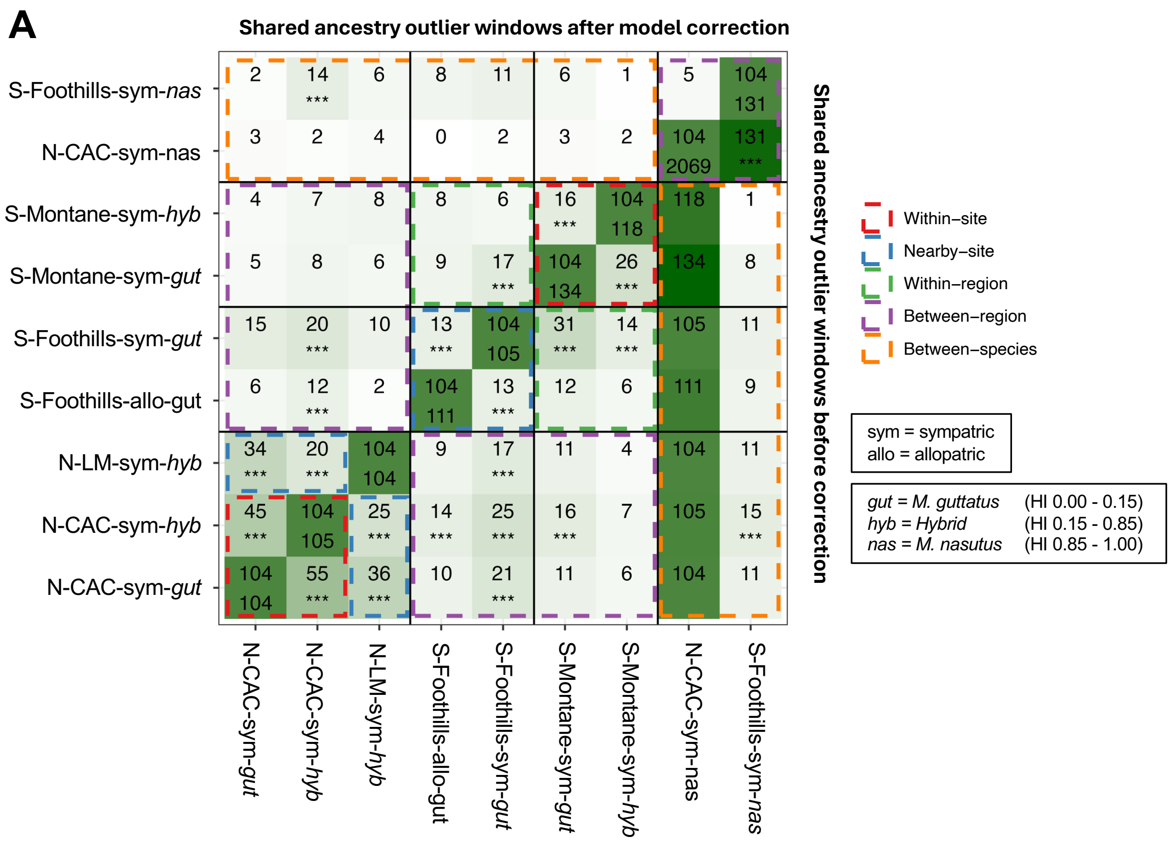

**Figure S6. Residual ancestry frequency distributions in reproductive isolation QTL regions, recalculated with reduced window dataset.** Compare to Main Text Figure 5. As in the full dataset, Northern hybrid groups show a significant reduction in residual ancestry frequency for N- QTL regions, but no significant increase in N+ QTL regions. The Southern Montane *hybrid* group and Northern CAC *nasutus* group do show significant increases in N+ QTL residual ancestry, consistent with the full dataset, though Southern Foothills allopatric and sympatric *guttatus*, which were both significantly increased in the full dataset, have no significant increase in ancestry frequency with the reduced dataset.

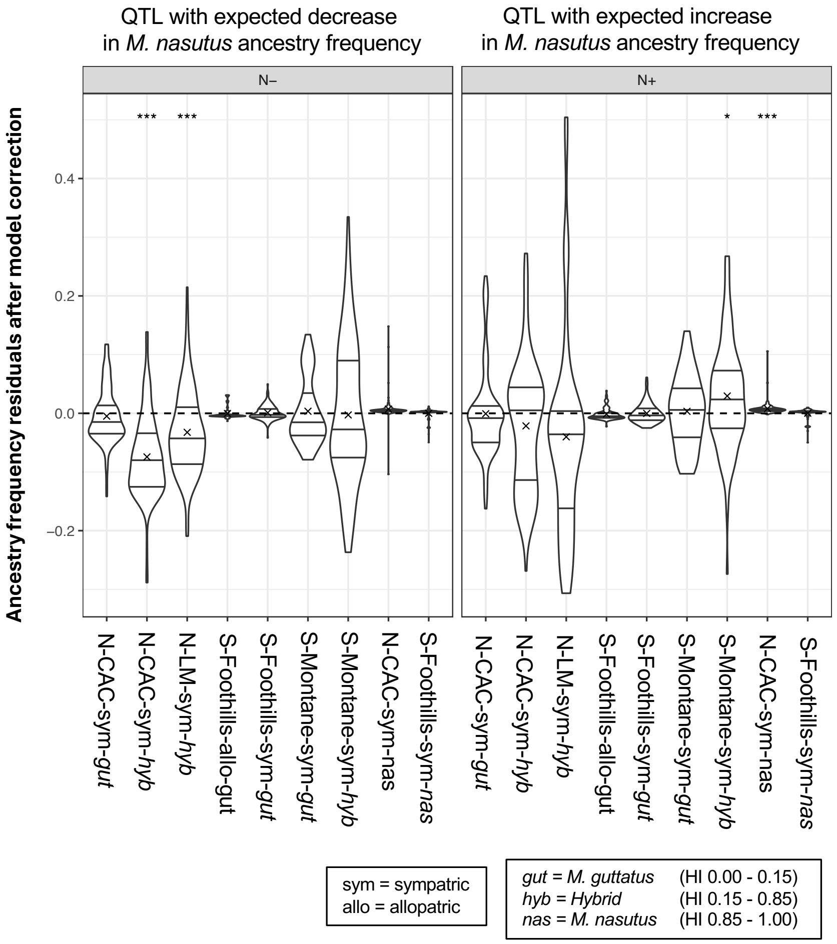

**Figure S7. Distance to the nearest sympatric location is negatively correlated with average hybrid index of allopatric Foothills locations.** Distance is given in meters, calculated using GPS coordinates in the R package 'geosphere'. The three allopatric locations with mean hybrid index >0.01 were designated as "allopatric-high" populations

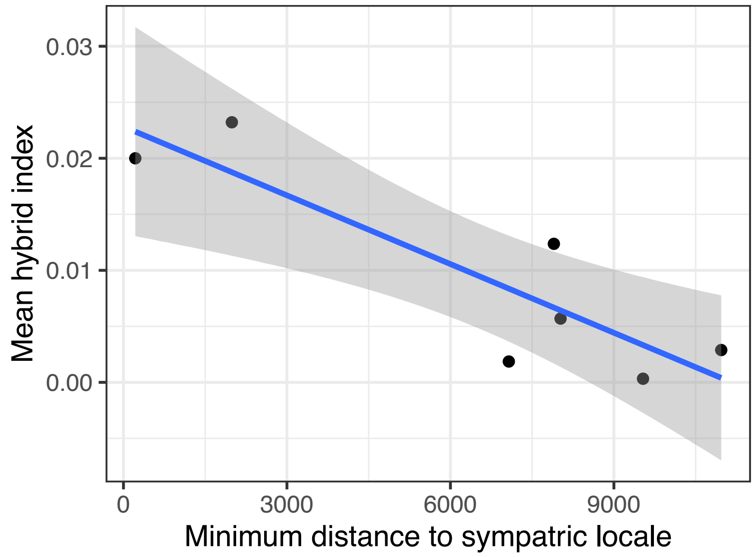

**Figure S8. Proportion of variance in ancestry frequency explained by genomic features, missingness, and other sample groups, with the Southern Foothills allopatric split into allopatric-low and allopatric-high subgroups.** Compare to Main Text Figure 3. The allopatric-high group consists of the three allopatric populations with higher average hybrid index (>0.01), while the allopatric-low group has lower average hybrid index (<0.01). The Allopatric-high subgroup had similar correlation patterns to the full allopatric group, suggesting that much of the allopatric signal of shared ancestry frequency patterns is driven by the allopatric populations with slightly higher total amounts of hybrid ancestry. However, correlations with other sample groups, particularly the Southern Foothills sympatric group, were still significant for the Allopatric-low subgroup, indicating that even these populations have some residual ancestry, likely derived from nearby sympatric populations via gene flow.

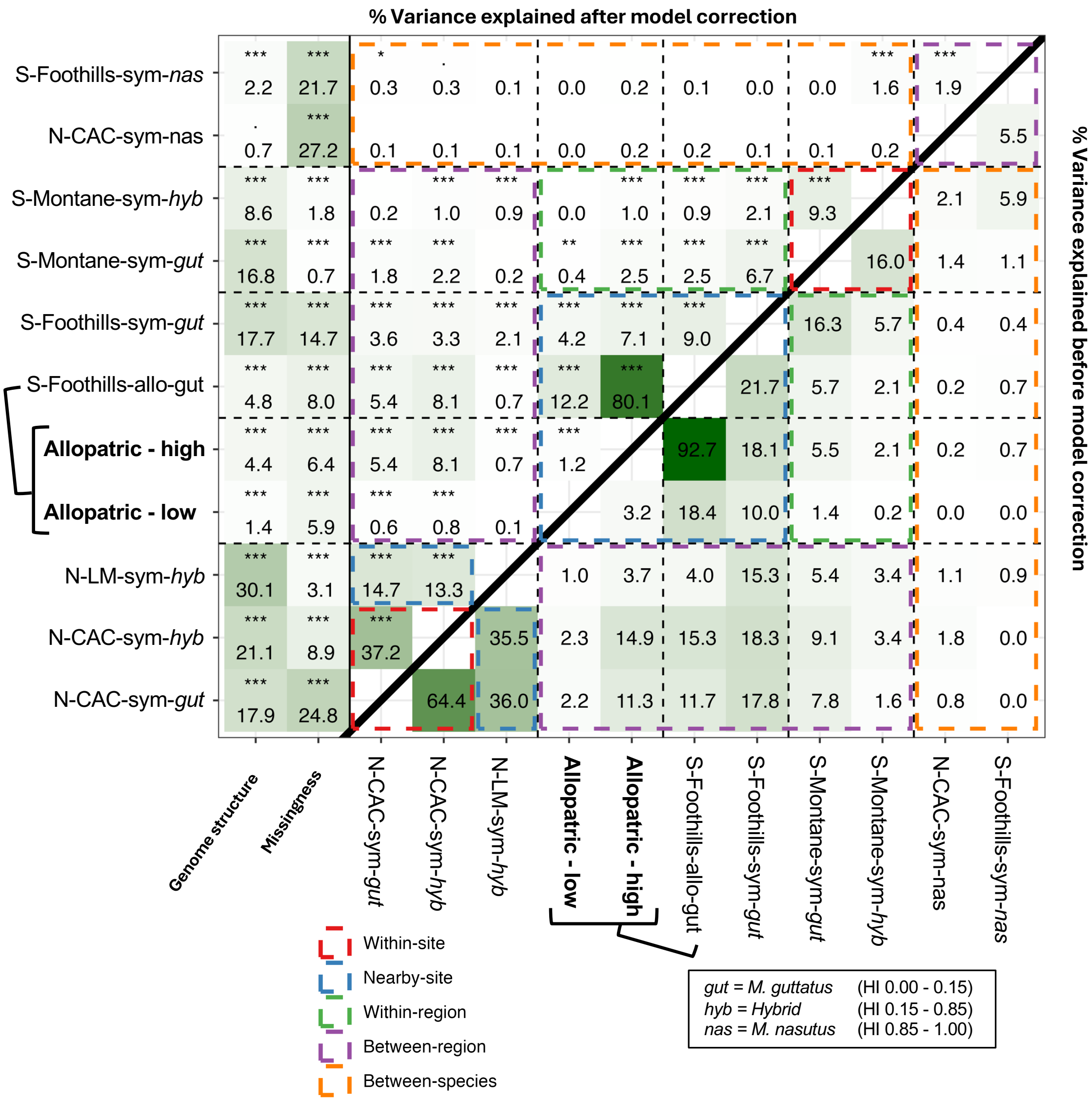

**Figure S9. Overlap among ancestry frequency outlier windows across groups, after correction for genomic features and missingness, with the Southern Foothills allopatric group split into allopatric-low and allopatric-high subgroups.** Compare to Main Text Figure 4A. The allopatric-high group consists of the three allopatric populations with higher average hybrid index (>0.01), while the allopatric-low group has lower average hybrid index (<0.01). The Allopatric-low group does not have significant overlap in outliers with Northern or Montane groups, but does share a significant number of overlapping outliers with the Southern Foothills sympatric group, consistent with gene flow from sympatry carrying a few introgressed alleles into all of our allopatric populations.

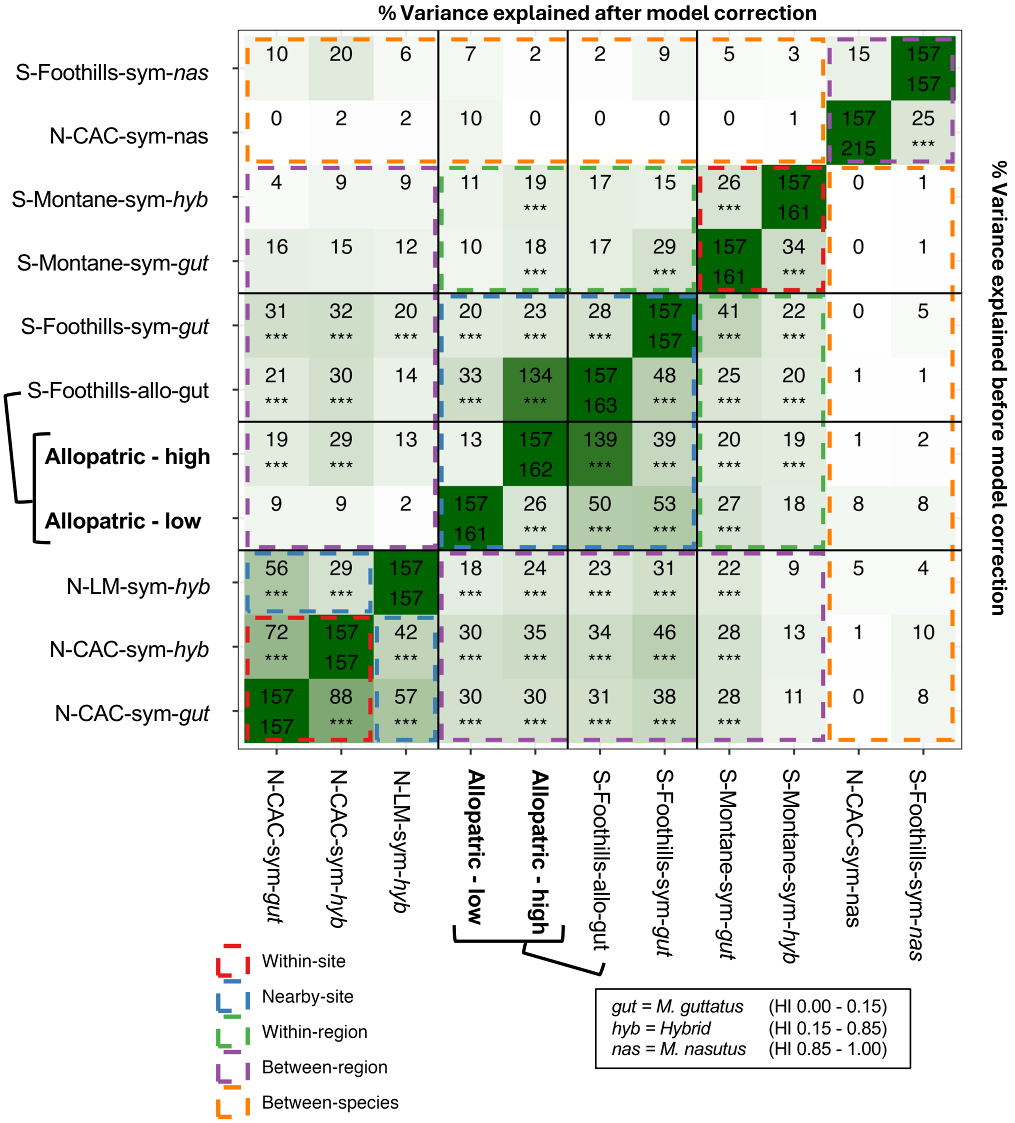

**Figure S10. Ancestry frequencies across all groups for chromosomes Chr01-Chr04.** Ancestry frequencies (colored points and grey line) vs. predictions from genomic structure and missingness modeling (black line) for sampling groups. Colors indicate the empirical quantile within the genome-wide distribution of residual ancestry frequency (i.e., the difference between observed ancestry frequencies and model predictions).

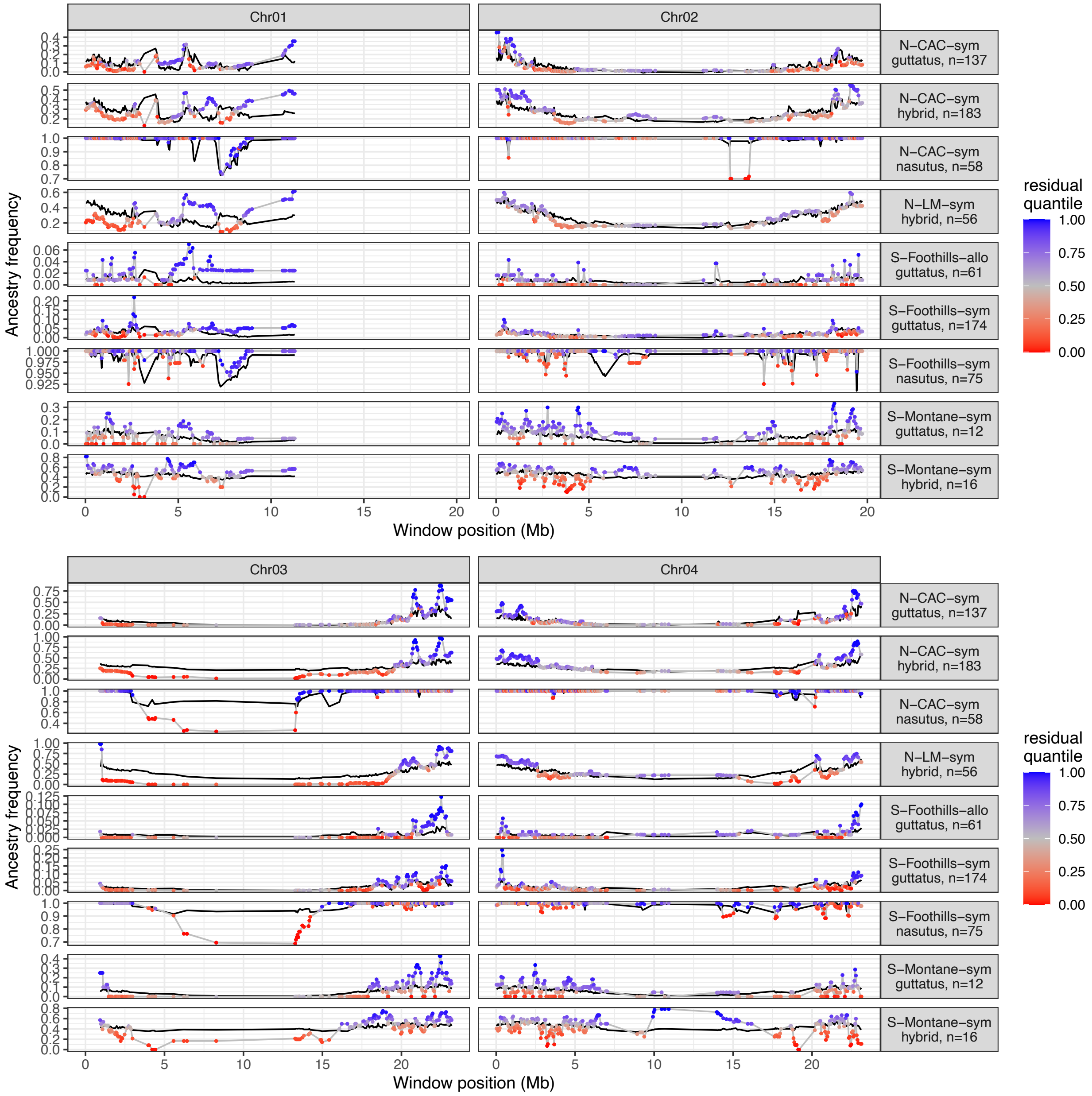

**Figure S11. Ancestry frequencies across all groups for chromosomes Chr05-Chr08.** Ancestry frequencies (colored points and grey line) vs. predictions from genomic structure and missingness modeling (black line) for sampling groups. Colors indicate the empirical quantile within the genome-wide distribution of residual ancestry frequency (i.e., the difference between observed ancestry frequencies and model predictions).

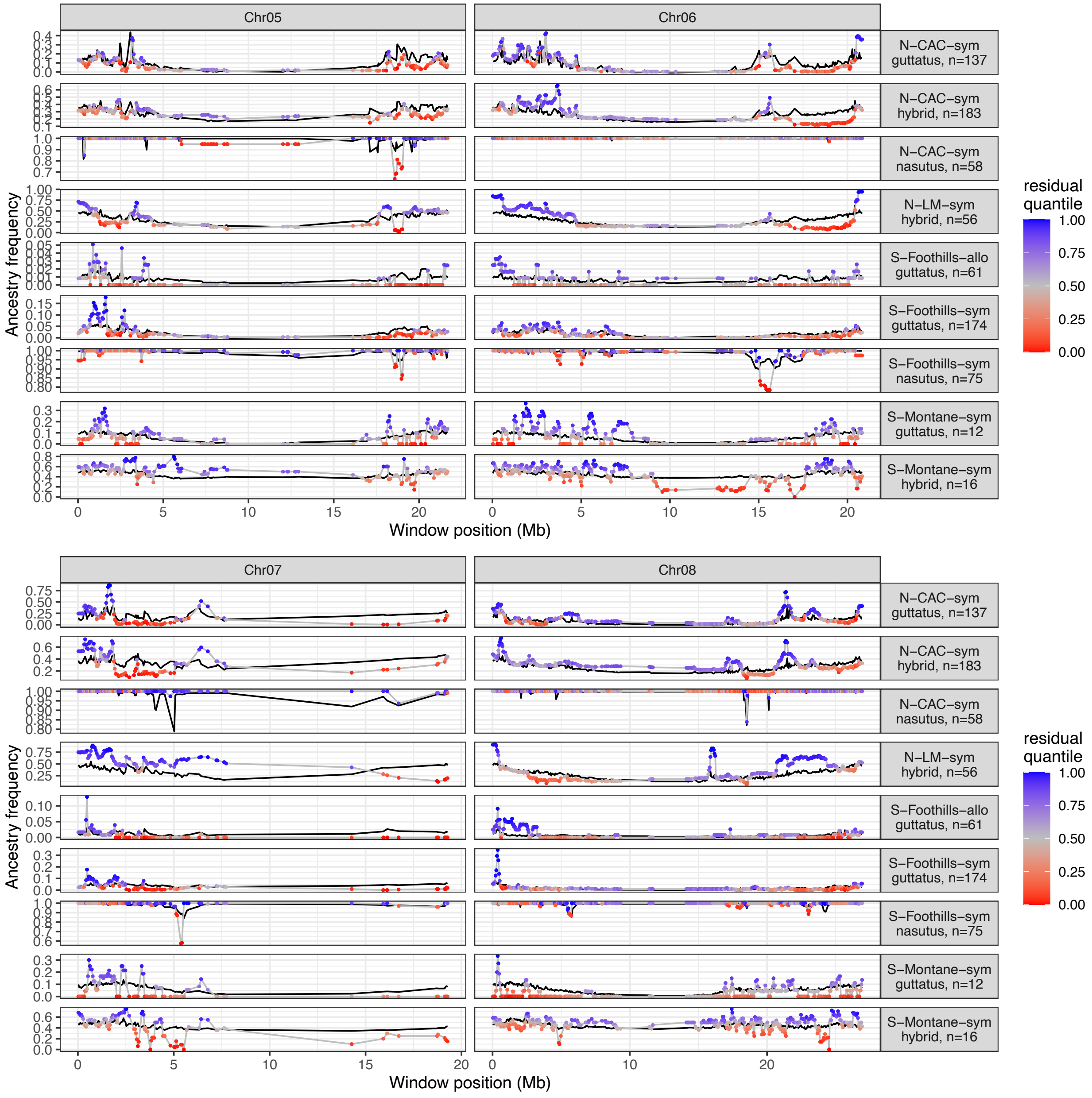

**Figure S12. Ancestry frequencies across all groups for chromosomes Chr09-Chr12.** Ancestry frequencies (colored points and grey line) vs. predictions from genomic structure and missingness modeling (black line) for sampling groups. Colors indicate the empirical quantile within the genome-wide distribution of residual ancestry frequency (i.e., the difference between observed ancestry frequencies and model predictions).

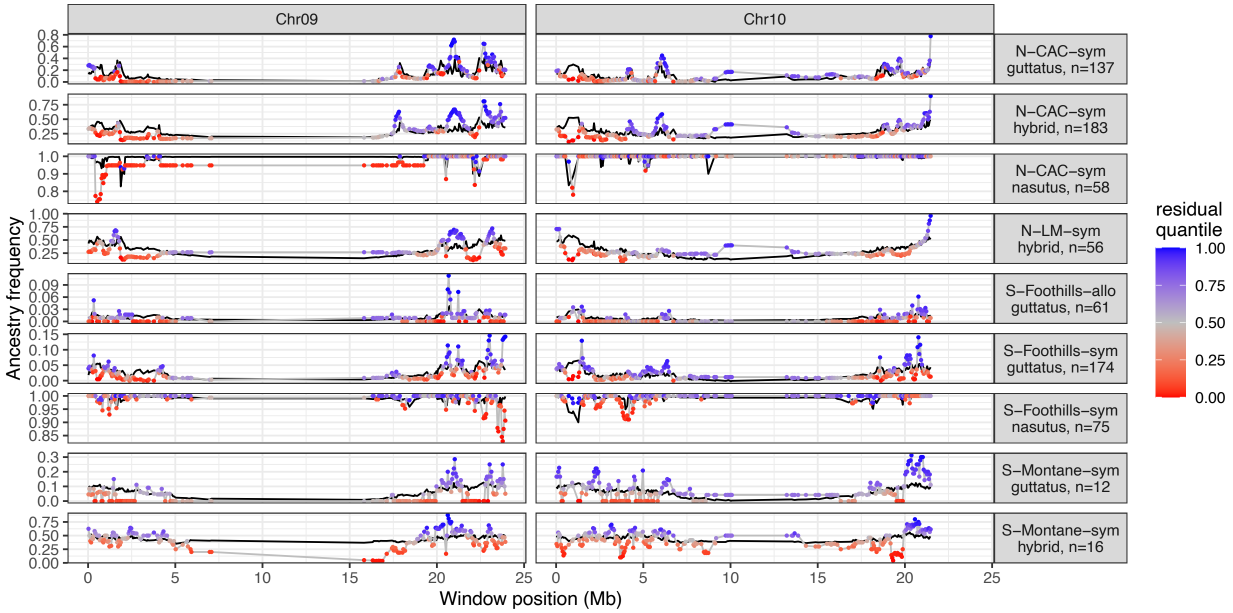


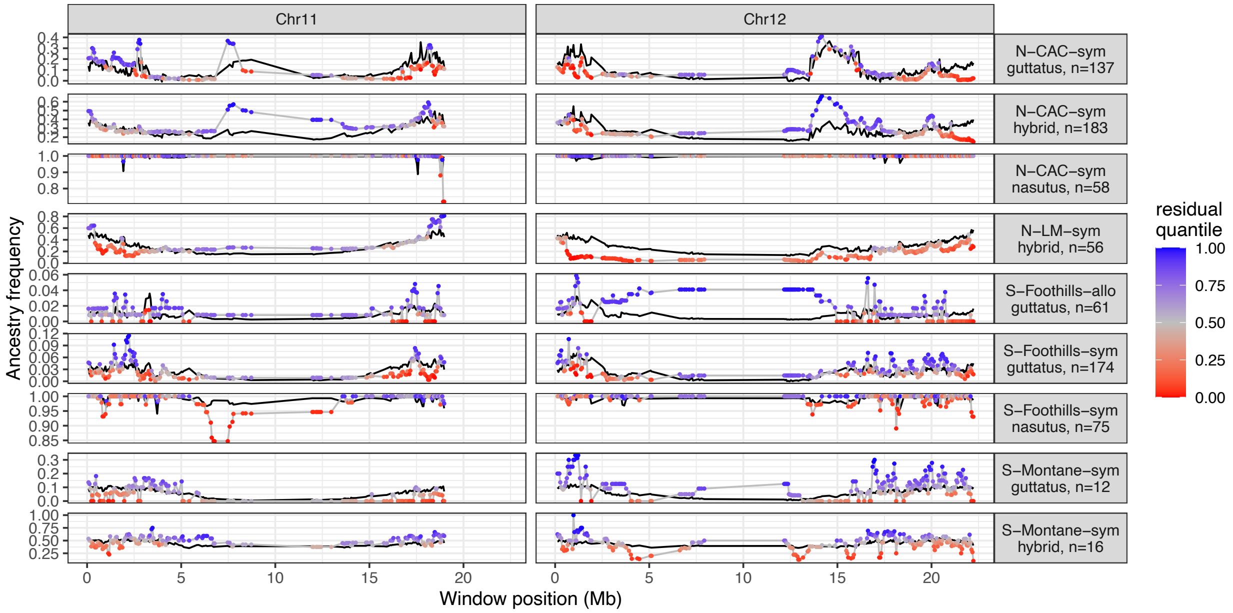

**Figure S13. Ancestry frequencies across all groups for chromosomes Chr13-Chr14.** Ancestry frequencies (colored points and grey line) vs. predictions from genomic structure and missingness modeling (black line) for sampling groups. Colors indicate the empirical quantile within the genome-wide distribution of residual ancestry frequency (i.e., the difference between observed ancestry frequencies and model predictions).

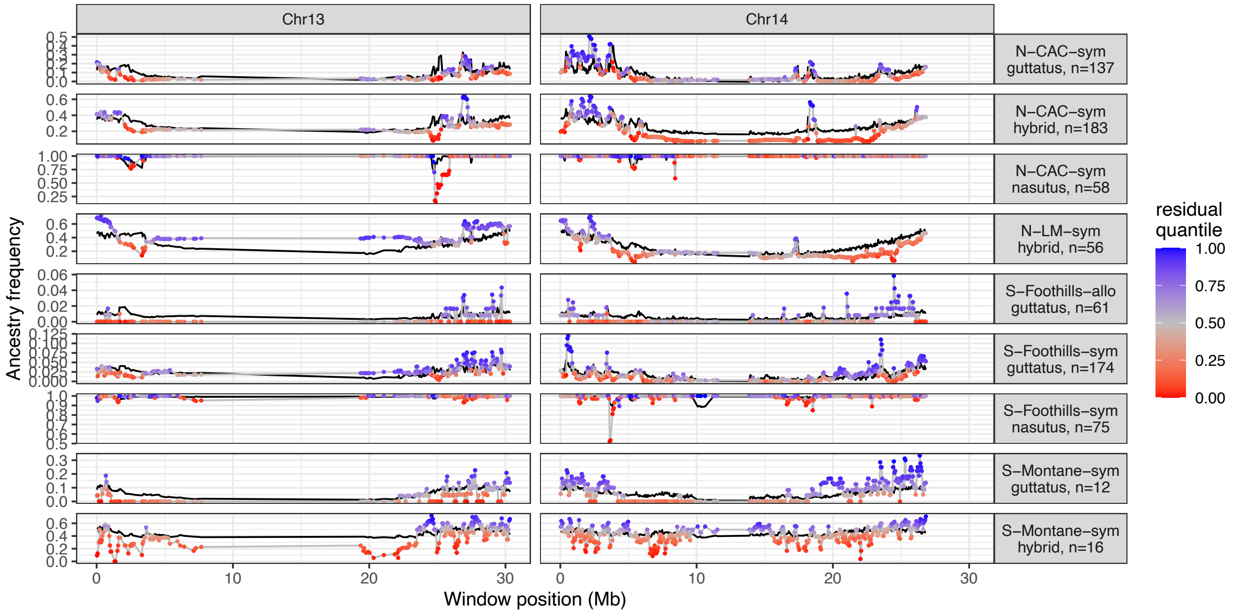
