## Supplementary Tables for "On the causes of correlated genomic ancestry across contrasting hybridization histories in a monkeyflower species pair"

These supplementary tables accompany the manuscript "On the causes of correlated genomic ancestry across contrasting hybridization histories in a monkeyflower species pair" by Farnitano, Matthew C., V. Alex Sotola, and Andrea L. Sweigart.

**Table S1. Sample sizes before and after filtering across all groups**

| **Pre-filtering** | *M. guttatus* HI<0.15 | *hybrid* 0.15<=HI<=0.85 | *M. nasutus* HI>0.85 | No data | *Totals* |
|---|---|---|---|---|---|
| Northern_CC_sympatric | 150 | 196 | 68 | 25 | 439 |
| Northern_LM_sympatric | 3 | 56 | 3 | 2 | 64 |
| Southern_foothills_allopatric | 61 | 0 | 0 | 3 | 64 |
| Southern_foothills_sympatric | 12 | 18 | 4 | 4 | 38 |
| Southern_montane_sympatric | 187 | 4 | 75 | 14 | 280 |
| *Totals* | 413 | 274 | 150 | 48 | 885 |

| **Post-filtering** | *M. guttatus* HI<0.15 | *hybrid* 0.15<=HI<=0.85 | *M. nasutus* HI>0.85 | *Totals* |
|---|---|---|---|---|
| Northern_CC_sympatric | 137 | 183 | 58 | 378 |
| Northern_LM_sympatric | 0** | 56 | 3 | 59 |
| Southern_foothills_allopatric | 61 | 0** | 0** | 61 |
| Southern_foothills_sympatric | 12 | 16 | 4** | 32 |
| Southern_montane_sympatric | 174 | 2** | 75 | 251 |
| *Totals* | 384 | 257 | 140 | 781 |

**categories with <10 individuals after filtering were not used for ancestry correlations, outlier identification, or other groupwise comparative analyses

**Table S2.** ***M. guttatus*** **and** ***M. nasutus*** **individuals used in an ancestry reference panel.**

| Line | Lat | Long | category | *M. nasutus* ancestry proportion | SRA accession |
|---|---|---|---|---|---|
| CAC415 | 45.71076 | -121.3667 | CC admixed *M. guttatus* | 0.0967 | SRX21547086 |
| CAC134 | 45.71076 | -121.3667 | CC admixed *M. guttatus* | 0.1014 | SRX21547082 |
| CAC141 | 45.71076 | -121.3667 | CC admixed *M. guttatus* | 0.1073 | SRX21547083 |
| CAC162 | 45.71076 | -121.3667 | CC admixed *M. guttatus* | 0.1135 | SRX21547084 |
| CACG6 | 45.71076 | -121.3667 | CC admixed *M. guttatus* | 0.1625 | SRX525044 |
| CAC262 | 45.71076 | -121.3667 | CC admixed *M. guttatus* | 0.2702 | SRX21547085 |
| CAC112 | 45.71076 | -121.3667 | CC admixed *M. guttatus* | 0.3127 | SRX21547081 |
| CAC110 | 45.71076 | -121.3667 | CC admixed *M. guttatus* | 0.3155 | SRX21547080 |
| NHN26 | 49.273 | -124.16 | *M. nasutus* | 0.9884 | SRX525051 |
| CACN9 | 45.71076 | -121.3667 | *M. nasutus* | 0.9974 | SRR1259271 |
| JGI_SF.sampled* | 45.264 | -121.022 | *M. nasutus* | 0.9987 | PRJNA1112462 |
| Koot | 48.104 | -115.983 | *M. nasutus* | 0.9989 | SRR1259272 |
| MEN104 | 37.828 | -120.344 | *M. nasutus* | 0.9994 | SRS2770656,7,8 |
| AHQT1G | 44.431 | -110.813 | Northern *M. gutttatus* | 0.0000 | SRX142379 |
| ATTU | 55.46 | -4.63 | Northern *M. gutttatus* | 0.0000 | SRX10011990 |
| JGI_IM62.sampled* | 44.400833 | -122.14917 | Northern *M. gutttatus* | 0.0000 | PRJNA1112459 |
| JGI_IM767.sampled* | 44.400833 | -122.14917 | Northern *M. gutttatus* | 0.0000 | PRJNA1112461 |
| MAR | 43.4786 | -123.29445 | Northern *M. gutttatus* | 0.0000 | SRX030542 |
| TSG3 | 53.418833 | -131.91573 | Northern *M. gutttatus* | 0.0000 | SRX2019854 |
| YJS6 | 44.9512 | -114.5845 | Northern *M. gutttatus* | 0.0008 | SRX030545 |
| BOG10 | 41.923611 | -118.80583 | Northern *M. gutttatus* | 0.0010 | SRX030570 |
| GUT5 | 42.71 | -118.57 | Northern *M. gutttatus* | 0.0083 | SRX10011991 |
| INV | 38.08 | -122.87 | Southern *M. guttatus* | 0.0000 | SRX6914899 |
| Odell_S11 | 43.54795 | -121.96278 | Southern *M. guttatus* | 0.0002 | SRR10194640 |
| SWB | 39.035983 | -123.69047 | Southern *M. guttatus* | 0.0008 | SRX030679 |
| SLP9 | 37.8482564 | -120.46187 | Southern *M. guttatus* | 0.0013 | SRX142377 |
| REM8G | 38.860433 | -122.41147 | Southern *M. guttatus* | 0.0014 | SRX030546 |
| PED5 | 32.982928 | -110.78417 | Southern *M. guttatus* | 0.0018 | SRR071969 |

| | | | | | |
|---|---|---|---|---|---|
| SCH | 39.018467 | -107.03527 | Southern *M. guttatus* | 0.0024 | SRX371892 |
| LMC24 | 38.863983 | -123.08392 | Southern *M. guttatus* | 0.0057 | SRX030680 |
| CSS4 | 38.861111 | -122.41528 | Southern *M. guttatus* | 0.0058 | SRX6435296 |
| SHG | 38.861111 | -122.41528 | Southern *M. guttatus* | 0.0084 | SRX10011994 |
| YVO6 | 37.723367 | -119.74643 | Southern *M. guttatus* | 0.0371 | SRX6914891 |

*reads from ultra-high-coverage JGI accessions were randomly downsampled to 60,000,000 per line prior to alignment.

**Table S3. Relationship between ancestry and genomic features before and after excluding pericentromeric regions.** Each value is the adjusted R^2 for the correlation between ancestry of a sample group and one genomic feature. Significance is based on an F-test with bonferroni-corrected (for 9 tests) p-values: *** = p<0.001, ** = p<0.01, * = p<0.05, . = p<0.1

| **All windows** | Relative chomosomal position | Genes per window | Recombination rate (1Mb) | Recombination rate (50kb) | Genes per cM |
|---|---|---|---|---|---|
| N-CAC-sym *guttatus* | 0.147*** | 0.074*** | 0.041*** | 0.013*** | 0 |
| N-CAC-sym *hybrid* | 0.176*** | 0.073*** | 0.051*** | 0.013*** | 0 |
| N-CAC-sym *nasutus* | 0.002 | 0.003** | 0.006*** | 0.003** | 0.001 |
| N-LM-sym *hybrid* | 0.248*** | 0.113*** | 0.074*** | 0.015*** | 0.008*** |
| S-Foothills-allo *guttatus* | 0.029*** | 0.014*** | 0.013*** | 0.002. | 0 |
| S-Montane-sym *guttatus* | 0.127*** | 0.099*** | 0.119*** | 0.054*** | 0 |
| S-Montane-sym *hybrid* | 0.037*** | 0.048*** | 0.074*** | 0.039*** | 0 |
| S-Foothills-sym *guttatus* | 0.141*** | 0.095*** | 0.072*** | 0.019*** | 0.001 |
| S-Foothills-sym *nasutus* | 0.011*** | 0.013*** | 0.019*** | 0.009*** | 0 |
| **Excluding pericentromeric†** | | | | | |
| N-CAC-sym *guttatus* | 0.138*** | 0.05*** | 0.019*** | 0.003* | 0.007*** |
| N-CAC-sym *hybrid* | 0.208*** | 0.073*** | 0.04*** | 0.004* | 0.01*** |
| N-CAC-sym *nasutus* | 0 | 0 | 0 | 0 | 0 |
| N-LM-sym *hybrid* | 0.259*** | 0.089*** | 0.03*** | 0.003. | 0.018*** |
| S-Foothills-allo *guttatus* | 0.101*** | 0.027*** | 0.034*** | 0.001 | 0.004** |
| S-Montane-sym *guttatus* | 0.069*** | 0.039*** | 0.043*** | 0.01*** | 0 |
| S-Montane-sym *hybrid* | 0.038*** | 0.021*** | 0.054*** | 0.022*** | 0 |
| S-Foothills-sym *guttatus* | 0.132*** | 0.062*** | 0.04*** | 0.001 | 0.013*** |
| S-Foothills-sym *nasutus* | 0.006*** | 0.005** | 0.009*** | 0.001 | 0 |
| †windows with recombination rate <1cM/Mb OR relative chromosomal position <0.25 were considered pericentromeric | | | | | |

**Table S4. Breakdown of M. guttatus (HI<0.15) individuals within Southern populations**

| group | population | Distance (m) to nearest sympatric location | mean hybrid index | n total | samples with HI>=0.01 n | samples with HI>=0.01 proportion |
|---|---|---|---|---|---|---|
| Southern_montane_sympatric | TUO | | 0.0766 | 4 | 4 | 1.00 |
| Southern_montane_sympatric | HHT | | 0.0734 | 7 | 5 | 0.71 |
| Southern_montane_sympatric | HHR | | 0.0041 | 1 | 0 | 0.00 |
| Southern_foothills_sympatric | NBR | | 0.0264 | 11 | 9 | 0.82 |
| Southern_foothills_sympatric | DPR | | 0.0257 | 94 | 85 | 0.90 |
| Southern_foothills_sympatric | DPI | | 0.0200 | 5 | 4 | 0.80 |
| Southern_foothills_sympatric | MFA | | 0.0190 | 3 | 2 | 0.67 |
| Southern_foothills_sympatric | MFR | | 0.0186 | 14 | 14 | 1.00 |
| Southern_foothills_sympatric | OPG | | 0.0177 | 5 | 4 | 0.80 |
| Southern_foothills_sympatric | MFB | | 0.0158 | 4 | 2 | 0.50 |
| Southern_foothills_sympatric | JAC | | 0.0144 | 18 | 12 | 0.67 |
| Southern_foothills_sympatric | CST | | 0.0133 | 2 | 1 | 0.50 |
| Southern_foothills_sympatric | NDP | | 0.0095 | 2 | 1 | 0.50 |
| Southern_foothills_sympatric | RCR | | 0.0081 | 12 | 3 | 0.25 |
| Southern_foothills_sympatric | GVP | | 0.0064 | 4 | 0 | 0.00 |
| Southern_foothills_allopatric | **MOC** | 1989 | 0.0232 | 10 | 5 | 0.50 |
| Southern_foothills_allopatric | **RCF** | 7899 | 0.0124 | 12 | 7 | 0.58 |
| Southern_foothills_allopatric | **BFR** | 8021 | 0.0057 | 10 | 1 | 0.10 |
| Southern_foothills_allopatric | COP† | 10969 | 0.0029 | 8 | 0 | 0.00 |
| Southern_foothills_allopatric | GCH† | 7072 | 0.0019 | 10 | 0 | 0.00 |
| Southern_foothills_allopatric | RHI† | 9534 | 0.0003 | 11 | 0 | 0.00 |

**Bolded** populations: grouped into "allopatric-high"

† populations: grouped into "allopatric-low"

**Table S5. Raw and residual ancestry frequency quantiles at four fine-mapped reproductive isolation loci.** Values in the top or bottom 5% quantile are bolded; values in the top 20% quantile are in green text; values in the bottom 20% quantile are in red text. Neighborhood mean refers to the mean across windows within 500kb of the focal window (11 total windows per locus).

| | | | | Raw ancestry frequency | | Residual ancestry frequency | | Neighborhood mean ancestry frequency | | Neighborhood mean residual ancestry frequency | |
|---|---|---|---|---|---|---|---|---|---|---|---|
| Gene name | chrom | windowend | group | value | quantile | value | quantile | value | quantile | value | quantile |
| FT-related | Chr07 | 3350000 | N-CC-sym gut | 0.0175 | 0.190 | **-0.137** | **0.034** | 0.039 | 0.349 | -0.141 | 0.054 |
| FT-related | Chr07 | 3350000 | N-CC-sym hyb | 0.1450 | 0.123 | **-0.229** | **0.009** | 0.191 | 0.250 | -0.181 | 0.076 |
| FT-related | Chr07 | 3350000 | N-CC-sym nas | **1.0000** | **1.000** | 0.002 | 0.331 | **1.000** | **1.000** | 0.008 | 0.662 |
| FT-related | Chr07 | 3350000 | N-LM-sym hyb | 0.3750 | 0.694 | -0.067 | 0.337 | 0.483 | 0.798 | 0.100 | 0.715 |
| FT-related | Chr07 | 3350000 | S-FH-allo gut | 0.0000 | 0.546 | **-0.013** | **0.048** | 0.002 | 0.593 | -0.010 | 0.215 |
| FT-related | Chr07 | 3350000 | S-FH-sym gut | 0.0258 | 0.691 | -0.024 | 0.062 | 0.017 | 0.396 | -0.025 | 0.150 |
| FT-related | Chr07 | 3350000 | S-FH-sym nas | **1.0000** | **1.000** | 0.026 | 0.960 | 0.995 | 0.630 | 0.007 | 0.660 |
| FT-related | Chr07 | 3350000 | S-MO-sym gut | 0.2500 | 0.981 | 0.158 | 0.973 | 0.045 | 0.504 | -0.040 | 0.270 |
| FT-related | Chr07 | 3350000 | S-MO-sym hyb | **0.6667** | **0.967** | 0.249 | 0.983 | 0.337 | 0.313 | -0.102 | 0.318 |
| SVP-cluster | Chr08 | 25250000 | N-CC-sym gut | 0.0484 | 0.433 | -0.078 | 0.116 | 0.067 | 0.517 | -0.050 | 0.254 |
| SVP-cluster | Chr08 | 25250000 | N-CC-sym hyb | 0.1761 | 0.184 | **-0.172** | **0.037** | 0.239 | 0.414 | -0.078 | 0.244 |
| SVP-cluster | Chr08 | 25250000 | N-CC-sym nas | **1.0000** | **1.000** | 0.005 | 0.793 | **1.000** | **1.000** | 0.004 | 0.657 |
| SVP-cluster | Chr08 | 25250000 | N-LM-sym hyb | 0.3438 | 0.644 | -0.062 | 0.347 | 0.411 | 0.732 | 0.020 | 0.574 |
| SVP-cluster | Chr08 | 25250000 | S-FH-allo gut | 0.0104 | 0.784 | -0.009 | 0.136 | 0.008 | 0.709 | -0.007 | 0.347 |
| SVP-cluster | Chr08 | 25250000 | S-FH-sym gut | 0.0189 | 0.569 | -0.014 | 0.173 | 0.015 | 0.464 | -0.018 | 0.157 |
| SVP-cluster | Chr08 | 25250000 | S-FH-sym nas | **1.0000** | **1.000** | 0.004 | 0.605 | **1.000** | **1.000** | 0.002 | 0.366 |
| SVP-cluster | Chr08 | 25250000 | S-MO-sym gut | 0.1429 | 0.859 | 0.056 | 0.820 | 0.080 | 0.652 | -0.006 | 0.501 |
| SVP-cluster | Chr08 | 25250000 | S-MO-sym hyb | 0.6500 | 0.949 | 0.195 | 0.946 | 0.589 | 0.805 | 0.115 | 0.757 |
| pTAC13 | Chr13 | 30300000 | N-CC-sym gut | 0.0852 | 0.593 | -0.078 | 0.116 | 0.097 | 0.632 | -0.070 | 0.154 |
| pTAC13 | Chr13 | 30300000 | N-CC-sym hyb | 0.2857 | 0.592 | -0.092 | 0.188 | 0.286 | 0.591 | -0.094 | 0.185 |
| pTAC13 | Chr13 | 30300000 | N-CC-sym nas | **1.0000** | **1.000** | 0.003 | 0.430 | **1.000** | **1.000** | 0.002 | 0.364 |
| pTAC13 | Chr13 | 30300000 | N-LM-sym hyb | 0.5648 | 0.876 | 0.049 | 0.673 | 0.449 | 0.755 | -0.059 | 0.401 |
| pTAC13 | Chr13 | 30300000 | S-FH-allo gut | 0.0000 | 0.546 | -0.012 | 0.052 | 0.000 | 0.546 | -0.012 | 0.060 |
| pTAC13 | Chr13 | 30300000 | S-FH-sym gut | 0.0394 | 0.833 | 0.002 | 0.643 | 0.036 | 0.800 | -0.001 | 0.544 |
| pTAC13 | Chr13 | 30300000 | S-FH-sym nas | **1.0000** | **1.000** | 0.006 | 0.701 | 0.996 | 0.922 | 0.003 | 0.561 |
| pTAC13 | Chr13 | 30300000 | S-MO-sym gut | 0.1364 | 0.847 | 0.055 | 0.814 | 0.127 | 0.800 | 0.035 | 0.680 |
| pTAC13 | Chr13 | 30300000 | S-MO-sym hyb | 0.6429 | 0.943 | **0.198** | **0.950** | 0.509 | 0.618 | 0.042 | 0.577 |
| pTAC14 | Chr14 | 15900000 | N-CC-sym gut | 0.0221 | 0.257 | 0.013 | 0.677 | 0.023 | 0.266 | 0.008 | 0.638 |
| pTAC14 | Chr14 | 15900000 | N-CC-sym hyb | 0.0929 | 0.060 | -0.072 | 0.255 | 0.092 | 0.058 | -0.077 | 0.236 |
| pTAC14 | Chr14 | 15900000 | N-CC-sym nas | **1.0000** | **1.000** | 0.001 | 0.111 | **1.000** | **1.000** | 0.002 | 0.178 |
| pTAC14 | Chr14 | 15900000 | N-LM-sym hyb | 0.1250 | 0.122 | -0.010 | 0.510 | 0.115 | 0.102 | -0.037 | 0.418 |
| pTAC14 | Chr14 | 15900000 | S-FH-allo gut | 0.0000 | 0.546 | -0.003 | 0.521 | 0.000 | 0.554 | -0.003 | 0.522 |
| pTAC14 | Chr14 | 15900000 | S-FH-sym gut | 0.0000 | 0.072 | -0.005 | 0.417 | 0.001 | 0.089 | -0.006 | 0.377 |
| pTAC14 | Chr14 | 15900000 | S-FH-sym nas | 0.9867 | 0.199 | -0.009 | 0.124 | 0.996 | 0.796 | 0.003 | 0.542 |
| pTAC14 | Chr14 | 15900000 | S-MO-sym gut | 0.0000 | 0.375 | -0.031 | 0.360 | 0.000 | 0.375 | -0.034 | 0.339 |
| pTAC14 | Chr14 | 15900000 | S-MO-sym hyb | 0.3000 | 0.146 | -0.116 | 0.183 | 0.394 | 0.350 | -0.038 | 0.386 |

**Table S6. Grouping of ancestry window outliers into contiguous stretches**

| Group | | sample_size | nwindows | nstretches |
|---|---|---|---|---|
| Northern_CAC_sympatric | *guttatus* | 137 | 157 | 60 |
| Northern_CAC_sympatric | *hybrid* | 183 | 157 | 65 |
| Northern_LM_sympatric | *hybrid* | 56 | 157 | 70 |
| Southern_Foothills_allopatric | *guttatus* | 61 | 157 | 68 |
| --subgroup allopatric-low | *guttatus* | 29 | 157 | 96 |
| --subgroup allopatric-high | *guttatus* | 32 | 157 | 66 |
| Southern_Foothills_sympatric | *guttatus* | 174 | 157 | 65 |
| Southern_Montane_sympatric | *guttatus* | 12 | 157 | 66 |
| Southern_Montane_sympatric | *hybrid* | 16 | 157 | 99 |
| Northern_CAC_sympatric | *nasutus* | 58 | 157 | 75 |
| Southern_Foothills_sympatric | *nasutus* | 75 | 157 | 85 |
